# A large scale systemic RNAi screen in the red flour beetle *Tribolium castaneum* identifies novel genes involved in insect muscle development

**DOI:** 10.1101/397117

**Authors:** Dorothea Schultheis, Matthias Weißkopf, Christoph Schaub, Salim Ansari, Van Anh Dao, Daniela Grossmann, Upalparna Majumdar, Muhammad Salim Hakeemi, Nicole Troelenberg, Tobias Richter, Christian Schmitt-Engel, Jonas Schwirz, Nadi Ströhlein, Matthias Teuscher, Gregor Bucher, Manfred Frasch

## Abstract

Although muscle development has been widely studied in *Drosophila melanogaster* there are still many gaps in our knowledge, and it is not known to which extent this knowledge can be transferred to other insects. To help in closing these gaps we participated in a large-scale RNAi screen that used the red flour beetle, *Tribolium castaneum*, as a screening platform. The effects of systemic RNAi were screened upon double-stranded RNA injections into appropriate muscle-EGFP tester strains. Injections into pupae were followed by the analysis of the late embryonic/early larval muscle patterns, and injections into larvae by the analysis of the adult thoracic muscle patterns. Herein we describe the results of the first-pass screens with pupal and larval injections, which covered ~8,500 and ~5,000 genes, respectively, of a total of ~16,500 genes of the *Tribolium* genome. Apart from many genes known from *Drosophila* as regulators of muscle development, a collection of genes previously unconnected to muscle development yielded phenotypes in larval body wall and leg muscles as well as in indirect flight muscles. We then present the main candidates from the pupal injection screen that remained after being processed through a series of verification and selection steps. Further, we discuss why distinct though overlapping sets of genes are revealed by the *Drosophila* and *Tribolium* screening approaches.

## Introduction

Muscle development in insects has been studied primarily in the dipteran *Drosophila melanogaster*, whereas much less is known about the regulatory mechanisms guiding muscle development in other insect orders. Thus, it is unknown whether more distantly-related insects, such as beetles, utilize largely the same processes and mechanisms to make muscles, or whether they differ in important aspects. Clearly, one shared process of holometabolous insects such as dipterans and coleopterans is that the musculature of an animal has to be developed twice; first the larval musculature during embryogenesis and second the adult musculature with completely different features during metamorphosis in the pupae.

In *Drosophila*, many of the genetic control mechanisms guiding the development of the multinucleated larval muscles of the body wall, also known as somatic muscles, have been uncovered during the past few decades. The earliest events in the hierarchy of zygotically active regulatory genes involve the activation of *twist* and *snail* in a ventral strip of cells in the blastoderm embryo, which are needed for the specification of the mesoderm and its invagination during gastrulation (Leptin 1991). *Tribolium castaneum* (*Tc*) *twist*, which is one of the few known regulators of mesoderm and muscle development in the red four beetle, fulfills similar early functions as *Drosophila twist* (Händel *et al.* 2005; Stappert *et al.* 2016). In *Drosophila* and likely in *Tribolium*, the subsequent spreading of the internalized mesoderm in tight contact with the overlying ectoderm is facilitated by FGF signals from the ectoderm (Wilson and Leptin 2000; Sharma *et al.* 2015). Subsequent patterning events that largely depend on signals from the ectoderm subdivide the mesoderm along the anterior-posterior and dorsoventral axis within each parasegmental unit, which leads to the formation of the anlagen giving rise the somatic, cardiac, and visceral muscles. (Baylies *et al.* 1995; Lee and Frasch 2000; Bodmer and Frasch 2010; Azpiazu *et al.* 1996; Baylies and Bate 1996; Riechmann *et al.* 1997; Lee and Frasch 2000). The somatic mesoderm then is subdivided further into domains that are competent to respond to localized and temporally-regulated receptor tyrosine kinase (RTK) signals. These are mediated by the *Drosophila* epidermal growth factor receptor (EGFR) or, alternatively, the fibroblast growth factor receptor Heartless (Htl) (Frasch 1999; Baylies and Michelson 2001; Carmena *et al.* 1998). The antagonistic actions between these RTK signaling activities and Delta/Notch signaling activities within these equivalence groups of cells ultimately result in the formation of two types of myoblast within each group. The first consists of a single muscle progenitor, in which the RTK signaling cascade remains active and the Notch signaling cascade is inactive. The second consists of several adjacent cells, in which the RTK signaling cascade is off and Notch activity is on. The muscle progenitor divides asymmetrically and typically gives rise to two muscle founder cells, each being programmed to form a single somatic muscle. The specific identity of each muscle founder is defined by the expression and functions of specific combinations of so-called muscle identity genes that generally encode various types of transcription factors (de Joussineau *et al.* 2012; Dobi *et al.* 2015). Conversely, within the adjacent cells lacking RTK activities, high Notch signaling activities induce the transcription factor encoding gene *lameduck* (*lmd*), which defines these cells as fusion-competent myoblasts that, *a priori*, are not committed to specific muscle fates (Duan *et al.* 2001; Ruiz-Gomez *et al.* 2002).

During the next important event, myoblast fusion, fusion-competent myoblasts fuse sequentially to each muscle founder cell and nascent myotube to generate a specific body wall muscle (Kim *et al.* 2015; Deng *et al.* 2017). The recognition and adhesion of the two types of myoblast occurs through the engagement of the immunoglobulin (Ig) domain proteins Sticks-and-stones (Sns) and Hibris (Hbs) on fusion-competent myoblasts with the related Ig domain proteins Kin of irre (Kirre) (aka, Dumbfounded, Duf) and Roughest (Rst, aka, IrreC) on the muscle founder cells. Downstream signaling cascades in both cell types lead to the differential assembly of polymerized actin structures at the prospective fusion site. Most prominently, actin-propelled protrusions from the fusion-competent myoblasts into the founder cells are thought to cause membrane rupture and fusion pores (Kim *et al.* 2015; Deng *et al.* 2017).

Towards the end and after myoblast fusion, the syncytial muscle precursors form extensions that migrate to the specific epidermal muscle attachment sites and make contacts with them. Several mechanisms regulating myotube guidance and the establishment of initial contacts have been identified (Schweitzer *et al.* 2010; Maartens and Brown 2015; Schulman *et al.* 2015). These include the release of Slit proteins from tendon cells, their binding to Robo receptors on the myotubes for proper guidance, subsequent arrest of migration upon the interaction of Robo with the Leucine-rich tendon-specific protein (Lrt) protein on the membranes of the tendon cells, as well as interactions between the trans-membrane protein Kon-tiki (Kon, aka Perdido), its cytoplasmic partner, Grip (a PDZ domain-containing protein), and the cell-surface protein, Echinoid (Ed). During muscle differentiation, myotendinous junctions and muscle-muscle connections at the attachment sites are stabilized by integrin-mediated adhesions to specific extracellular matrix structures between these cells, as well as by the intracellular linkage of integrin-associated proteins to the cytoskeleton in both muscle and tendon cells (Schnorrer and Dickson 2004; Schweitzer *et al.* 2010; Maartens and Brown 2015).

Muscle differentiation culminates in the assembly of the sarcomeric apparatus, proper positioning of the myonuclei, and the establishment of neuromuscular junctions (Nose 2012; Volk 2013; Schulman *et al.* 2015; Lemke and Schnorrer 2017). Two key regulators known to act in muscle differentiation in vertebrates, *MyoD* and *Mef2*, are present in *Drosophila* as single orthologs and regulate muscle differentiation (Michelson *et al.* 1990; Paterson *et al.* 1991; Bour *et al.* 1995; Lilly *et al.* 1995; Arredondo *et al.* 2001). *nautilus* (*nau*; *Drosophila MyoD*) is required for the formation and differentiation of a specific subset of larval muscles (Balagopalan *et al.* 2001). *Drosophila Mef2*, in addition to functioning in terminal muscle differentiation, has also an essential earlier role in myoblast fusion, likely via transcriptionally activating certain myoblast fusion genes (Sandmann *et al.* 2006; Brunetti *et al.* 2015).

Prior to adult muscle development, the vast majority of the larval body wall muscles are histolyzed in early pupae and the adult musculature is built mostly from stem cell-like cells that have been set aside during embryogenesis and are called adult muscle precursors (AMPs) (Gunage *et al.* 2017). Many, but not all of the known regulators of larval muscle development are being reutilized during adult muscle development (Gunage *et al.* 2017). Thoracic AMPs associated with wing and leg discs are being patterned by signals from the epidermal cell layer and thus assume differential muscle fates, such as direct versus indirect flight muscles in case of the wing disc (Sudarsan *et al.* 2001). The indirect flight muscles differ in their ultrastructure from all other fly muscles in that they exhibit fibrillar, stretch-activated myofibers instead of tubular myofiber organizations. The involvement of the transcription factor Spalt major (Salm) as a master regulator in fibrillar muscle development in the indirect flight muscles has been one of the few documented examples of conserved regulatory processes during muscle development between *Drosophila* and *Tribolium* (Schönbauer *et al.* 2011).

Large-scale loss-of-function screens have been performed by RNA interference (RNAi) and classical mutagenesis to fill the remaining gaps in our knowledge on the regulation of *Drosophila* muscle development (Schnorrer *et al.* 2010; Johnson *et al.* 2013; Hollfelder *et al.* 2014; Camuglia *et al.* 2018). Because each of these methods has its own limitations, such as the delayed action of inducible RNAi in *Drosophila* embryos and the lack of phenotypes with functionally-redundant genes, we chose to undertake an alternative approach. To identify new components required for normal development of the body wall (somatic) musculature in insects, and also to begin to understand the similarities and differences between dipterans and coleopterans in this process, we participated in a large-scale systemic RNAi screen in the red flour beetle *Tribolium castaneum*, termed “iBeetle” (Schmitt-Engel *et al.* 2015). Our general strategy was to identify genes with interesting knock-down phenotypes in the somatic musculature in *Tribolium* and subsequently study the functions of their orthologs in *Drosophila* in more detail. Herein we describe an overview of this screen and provide a first description of the obtained muscle phenotypes, and in the accompanying paper (Schultheis *et al.* 2019) we provide a detailed analysis of one of the identified genes, *Nostrin*, in *Drosophila*. In the current study, we describe examples of genes from the screen that are orthologous to known regulators of muscle development in *Drosophila* and, notably, the identification and verification of genes in the *Tribolium* screen that have not been implicated in muscle development in previous research.

## Materials and Methods

### *Tribolium* strains

All beetles were kept under standard conditions (Brown *et al.* 2009) on white wheat flour containing 5% dry yeast at 25 °C and shifted to 32 °C for the experiments. The following *Tribolium castaneum* stocks were used in this study: *San Bernardino (SB)*, *black* (Sokoloff *et al.* 1960), *piggyBac pig-19 (pBA19)* (Lorenzen *et al.* 2003), *D17Xred* (Schmitt-Engel *et al.* 2015).

### First-pass screens and rescreens for verification of phenotypes

Selection of targeted genes in the first-pass screens: For the first round of screening no selection of the targeted genes was done, although at the beginning of the screen there was a slight innate bias towards more highly expressed genes. For the second screening round, genes to be targeted by dsRNA fragments were selected if they fulfilled at least one of the following criteria: 1. Gene is homologous to a *D. melanogaster* protein of unknown function. 2. Gene product is conserved in *Drosophila* and others (significant blast hit in UniProt-SWISS-PROT-invertebrates database). 3. Gene has ortholog in other species but not in *Drosophila*. 4. Gene product is associated with GO terms of “development/transcription/signaling” (i.e., “molecular transducer activity”, “morphogen activity”, “nucleic acid binding transcription factor activity”, “protein binding transcription factor activity”, “translation regulator activity”, “receptor activity”, “receptor regulator activity”, “developmental process”, or their child terms). For additional information, see Schmitt-Engel *et al.* (2015).

#### Rescreens

The first rescreen was performed in *pig-19* and pupal injections of the original iB dsRNAs. Because of the aim to identify new *Drosophila* genes with functions in myogenesis the genes that have *Drosophila* orthologs with known roles in myogenesis and those that lack *Drosophila* orthologs were omitted (except for a few with striking and highly penetrant phenotypes). In addition, iB dsRNAs that produced phenotypes with very low penetrance were omitted, as were those that upon closer inspection of the database were likely to yield indirect effects on muscle development. In the second rescreen, the original iB dsRNAs as well as new dsRNAs with sequences that did not overlap with the original ones were injected into female pupae from the San Bernardino (SB) strain, and the muscle patterns were analyzed in embryos from a cross of these females with *pig-19* males.

### Cloning of *Tc-Mef2* and *Tc-duf/Kirre*

To synthesize first strand cDNA the Omniscript RT kit (Qiagen) was used. 5 µg total RNA derived from early embryonic stages were reverse transcribed utilizing oligo(dT) primers and following the manufacturer’s instructions. 1 µl of the cDNA synthesis reaction was subsequently used to amplify 1kb fragments of *Tc-Mef2* (*TC010850*) and *Tc-duf* (*TC002914*) by PCR using gene specific primers (*Tc-Mef2-F GTTTGATCGGTCCGTGCTAT*; *Tc-mef2-R GACCGCTCCAGGATATTGAA*; *Tc-duf-F ACGCGACCAGGAAATATCAC*; *Tc-duf-R GGAAGCTTGGTTCGGTGTAA*). The ~1kb *Tc-duf* fragment fully includes the iB_03469 sequences at its 3’ portion. The amplified PCR fragments were gel purified and cloned into the pCR©II-TOPO© vector using the TOPO©TA Cloning© Dual Promoter kit (ThermoFisher Scientific) following the manufacturer’s instructions.

### *Tc-mef2* and *Tc-Duf* RNA probe synthesis

To synthesize antisense DIG-labeled Riboprobes, 1 µg of linearized pCR©II-*Tc-mef2* or pCR©II-*Tc-duf*/*kirre* was *in vitro* transcribed utilizing the DIG RNA labelling kit (Roche) following the manufacturer’s instructions. The DIG-labelled RNA was purified using the RNA cleanup protocol of the RNeasy Kit (Qiagen). *Tribolium* fixation and *in situ* hybridization were performed as described previously (Tautz and Pfeifle 1989; Patel *et al.* 1994).

### Double-stranded RNA preparation and injections for RNAi

To synthesize dsRNA of *Tc-mef2* and *Tc-Duf*, 1 ng of pCR©II-*Tc-mef2* or pCR©II-*Tc-Duf*/*Kirre* were used in a PCR reaction using T7 and T7-SP6 primers. The amplified fragments were purified using the QIAquick Gel Purification Kit (Qiagen) and dsRNA was produced as described in Bucher *et al.* (2002). To induce parental RNAi 1 µg/µl of *Tc-mef2* or *Tc-Duf* dsRNA were injected into adult females as described in van der Zee *et al.* (2006).

The procedure of the iBeetle larval and pupal RNAi injection screen and the procedure for the analysis of late embryonic/early larval muscles are described in detail in Schmitt-Engel *et al.* (2015). The dsRNAs used in the iBeetle Screen were obtained from Eupheria Biotec GmbH (Dresden). To generate the dsRNAs for the rescreen cDNA was generated using the Transcriptor First Strand cDNA Synthesis Kit (Roche). The cDNA was then used to amplify fragments by PCR with the same primers as used by Eupheria Biotec GmbH (Dresden) for the original iBeetle Screen. The sequences of the dsRNA fragments in the primary screen (iB dsRNAs; annotated in the format iB_nnnnn) and of the dsRNA fragments non-overlapping with the iB fragments (annotated as iB_nnnnn_2) are accessible in http://ibeetle-base.uni-goettingen.de/gb2/gbrowse/tribolium/ (select track “iB dsRNA”). These dsRNAs were synthesized from PCR products using the MEGAscript^TM^ T7 Transcription Kit (Ambion).

### Microscopy

Live images from the screens and control animals were taken on a Leica M165 FC fluorescence dissecting microscope with a ProgRes C14 CCD camera (Jenoptic, Jena). The confocal image of a live pupal thorax was acquired on a Leica SP5II confocal laser scanning microscope with a 10x HC PL APO 0.40 CS objective using the LAS AF (Leica) software.

### Research materials and data availability

Materials produced in this study are available upon request. The authors affirm that all data necessary for confirming the conclusions of this article are represented fully within the article and its tables and figures with the exception of sequence information (e.g., for amplification primers) that is available at http://ibeetle-base.uni-goettingen.de/gb2/gbrowse/tribolium/.

## Results

### Larval and adult thoracic somatic muscle patterns in *Tribolium*

The RNAi screen took advantage of the systemic nature of parental RNA interference in *Tribolium castaneum* (Bucher *et al.* 2002). For screening we employed two different enhancer trap lines that expressed EGFP in muscles. The *pig-19* line carries a piggyBac insertion in the 3’ UTR of *TC003326* (*Actin-87E-like*) and expresses EGFP in all larval somatic and visceral muscles (Lorenzen *et al.* 2003). *pig-19* was used for analyzing RNAi phenotypes in the somatic musculature of late embryonic and first instar larvae upon double-stranded (ds) RNA injections into pupae. The second strain, *D17* (with unknown genomic insertion site), expresses EGFP in the indirect flight muscles in the thorax of late pupae and thus was used to score thoracic muscle patterns upon dsRNA injections into larvae.

Before screening we used these expression patterns to characterize the wild type pattern of the larval body wall musculature and the late pupal thoracic musculature in more detail. As shown with *pig-19*-EGFP, each abdominal segment displays a stereotypical arrangement of ca. 29 syncytial muscle fibers underneath the body wall, which exclude ventral areas where the CNS is located and the dorsal midline where the unlabeled dorsal vessel is positioned (Fig. 1A - E). In the thoracic segments, which unlike *Drosophila* embryos carry appendices, a modified muscle pattern is observed (Fig. 1A - D) and a stereotypic muscle pattern is also seen in the legs (Fig. 1F). The muscle arrangement in each abdominal segment into dorsal, lateral, and ventral groups, as well as their orientations as longitudinal, oblique, transverse, and acute muscles within these groups, are strongly reminiscent of the well-characterized pattern of abdominal body wall muscles in *Drosophila* (Fig. 1G - I) (Bate 1993). However, the details differ and due to the current lack of conserved markers for individual muscles or sets of muscles it is presently unclear whether any of these muscles are homologous between the two insect species.

**Figure 1.**
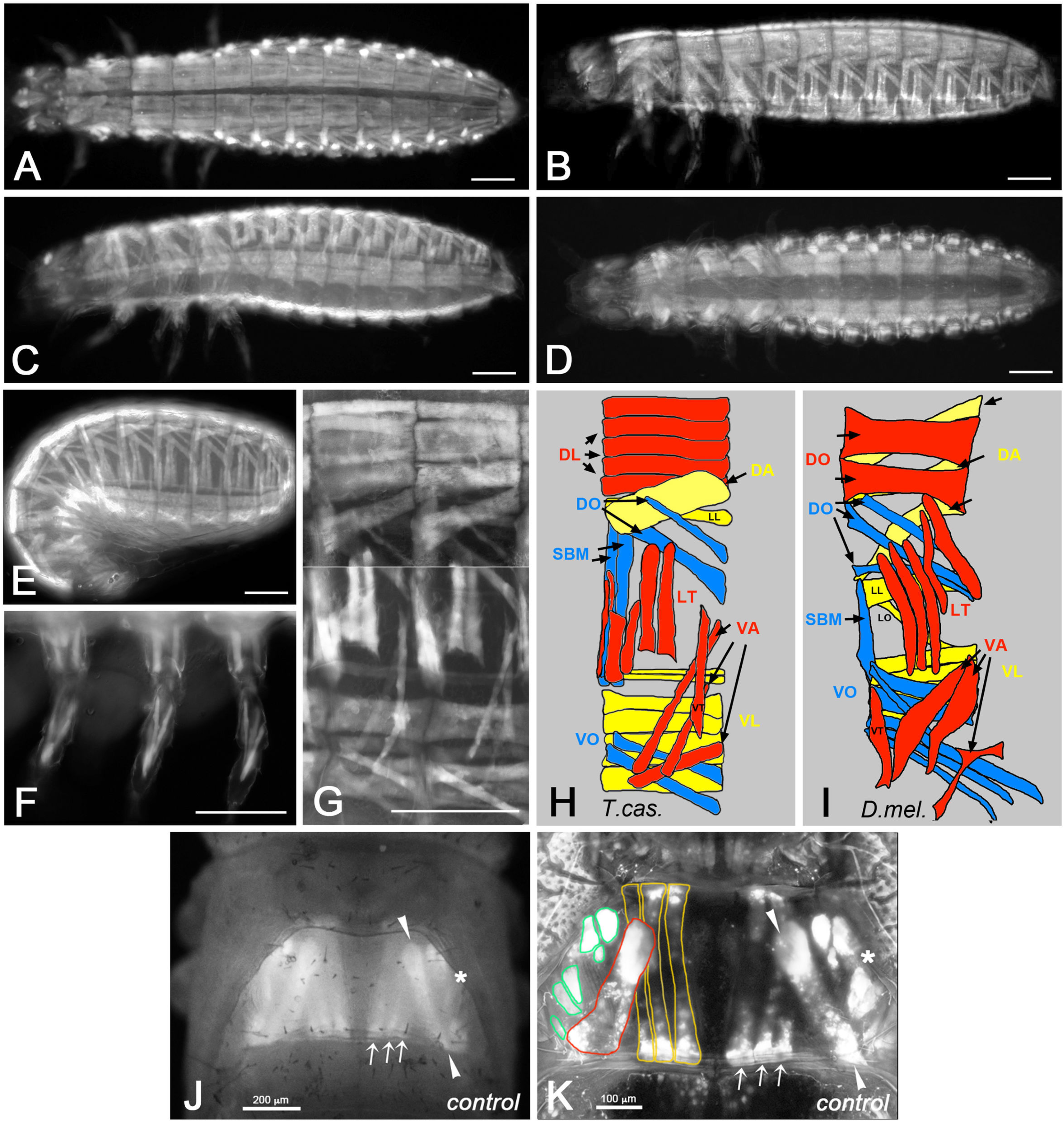
Somatic muscle pattern in *Tribolium castaneum* and *Tribolium* orthologs of known *Drosophila* myogenesis regulators. **(A)** to **(G)** show *Tribolium castaneum pig-19* enhancer trap 1^st^ instar larvae and embryos imaged live for EGFP expression in the somatic musculature. **(A)** Newly hatched 1^st^ instar larva, view of dorsal muscle pattern. **(B)** Newly hatched 1^st^ instar larva, view of dorsal-lateral muscle pattern. **(C)** Newly hatched 1^st^ instar larva, view of ventral-lateral muscle pattern. **(D)** Newly hatched 1^st^ instar larva, view of ventral muscle pattern. **(E)** Late stage embryo prior to hatching, view of lateral muscle pattern. **(F)** View of 1^st^ instar larval leg muscle pattern. **(G)** Newly hatched 1^st^ instar larva, high magnification view of muscle pattern in two abdominal segments (composite of a dorsal-lateral and a ventral-lateral view from two different animals, separated by white line). **(H)** Schematic representation of late embryonic muscle pattern in an abdominal segment from *Tribolium castaneum*. Note that the external-to-internal orders of the muscles and the exact numbers of DL and VL muscles are tentative. **(I)** Schematic representation of late embryonic muscle pattern in an abdominal segment from *Drosophila* (abbreviations: DA: dorsal acute; DL: dorsal longitudinal; DO: dorsal oblique; LT: longitudinal transverse; SBM: segment border muscle; VA: ventral acute; VL: ventral longitudinal; VO: ventral oblique muscles; for nomenclature see (Bate 1993)). **(J)** Thorax of *Tribolium castaneum D17* enhancer trap at late pupal stage. Shown is a dorsal view of indirect flight muscles taken with a fluorescence dissecting microscope as used in the screen. On either side, three dorsal longitudinal flight muscles (arrows), one dorsal oblique flight muscle (arrowheads), and a lateral area with unresolved dorsoventral flight muscles (asterisk) are marked by EGFP in the second thoracic segment. **(K)** Second thoracic segment of *D17* pupa as in (J) imaged by confocal microscopy. In addition to the dorsal longitudinal (arrows and yellow outlines) and dorsal oblique muscles (arrowheads and red outline), about six dorsoventral flight muscles (asterisk and green outlines) can be distinguished in lateral areas. Scale bars: A - G, K: 100μm; J: 200 μm.

*D17*-EGFP specifically marks the indirect flight muscles in the second thoracic segment starting from late pupal stages. In dorsal views of pupae observed under a fluorescence stereo microscope as used in the larval injection screen, three thin dorsal longitudinal flight muscles arranged medially in parallel and a broader dorsal oblique flight muscle inserted more laterally at its posterior can be discerned in each hemithorax (Fig. 1J). In lateral areas, the dorsoventral flight muscles can be seen. Under screening conditions these were evaluated as a single entity because they could not be resolved as individual muscles under the dissecting microscope. When examined by confocal microscopy, optical sections of at least six dorsoventral flight muscles can be recognized on either side, in addition to the dorsal longitudinal and dorsal oblique muscles also seen in the dissecting microscope (Fig. 1K).

### Positive controls with *Tribolium* orthologs of *Drosophila* myogenic regulators

Before embarking on the RNAi screen we sought evidence for similarities in muscle development between the two species of insect and tested whether RNAi-induced muscle phenotypes can be obtained for *Tribolium castaneum* (*Tc*) orthologs of myogenic regulators in *Drosophila*. The expression of the early mesodermal regulator Twist in the embryonic mesoderm of *Tribolium* has been documented extensively (Händel *et al.* 2005; Stappert *et al.* 2016), and as expected, RNAi against *Tc-twist* in *pig-19* led to a complete absence of muscles (data not shown, see http://ibeetle-base.uni-goettingen.de). *TC002914*, the single ortholog of *Drosophila kirre* (aka *duf*) and *roughest* (*rst*), which in *Drosophila* are essential for myoblast fusion in a functionally redundant manner, is expressed in somatic mesodermal cells of the body wall and limbs in *Tribolium* embryos (Fig. 2A). Whereas *Drosophila kirre* is expressed only in muscle founder cells, its paralog *rst* is expressed in both founder and fusion-competent myoblasts (Ruiz-Gomez *et al.* 2000; Strunkelnberg *et al.* 2001). The seemingly broader expression of *TC002914* as compared to *Drosophila kirre* may suggest that the expression of *TC002914* is more akin to that of *rst* in *Drosophila*. This interpretation is supported by the expression of *Tc-sticks-and-stones* (Tc-*sns*) (*TC032336*), which is similar albeit slightly narrower in the somatic mesoderm as compared to *Tc-kirre*/*rst* (data not shown). Potentially, *Tc-sns mRNA* is restricted to fusion-competent myoblasts like *Drosophila* Sns, which regulates myoblast fusion upon interaction with Kirre. Importantly, RNAi knock-down upon injections of *Tc-kirre*/*rst* dsRNA into in *pig-19* adult females led to strong reductions in both numbers and sizes of EGFP-stained muscles (Fig. 2B, B’), which corresponds to analogous phenotypes in *kirre rst* double mutants in *Drosophila* (Ruiz-Gomez *et al.* 2000; Strunkelnberg *et al.* 2001). In the most severe examples, only few small muscle fibers were present (Fig. 2B), whereas in milder cases presumably resembling partial knock-downs a fraction of the muscles were present but many others were very thin or missing (Fig. 2B’). Incidentally, this example also illustrates that RNAi screens in *Tribolium* have the potential to identify myogenic regulators that may have been missed in *Drosophila* screens due to the presence of functionally-redundant paralogs, if these have only a single ortholog in the beetle. Another example of similarities in myogenic regulation between the two species of insect is provided by *Mef2*, which in *Drosophila* is expressed in all muscle progenitors (and muscles) and encodes a crucial muscle differentiation factor (Lilly *et al.* 1994; Nguyen *et al.* 1994). In *Tribolium* embryos, the *Mef2* ortholog *TC010850* is also expressed broadly in the somatic mesoderm of the body wall and the limb (Fig. 2C). Like with *Drosophila Mef2* mutant embryos, knock-down of *TC010850* upon dsRNA injections into adult females caused almost complete loss of muscles as detected by EGFP in *pig-19* embryos (Fig 2D, D’) (Bour *et al.* 1995; Lilly *et al.* 1995). Additional examples of phenotypes of genes orthologous to known regulators of *Drosophila* myoblast fusion are shown in the accompanying paper (Schultheis *et al.* 2019).

**Figure 2.**
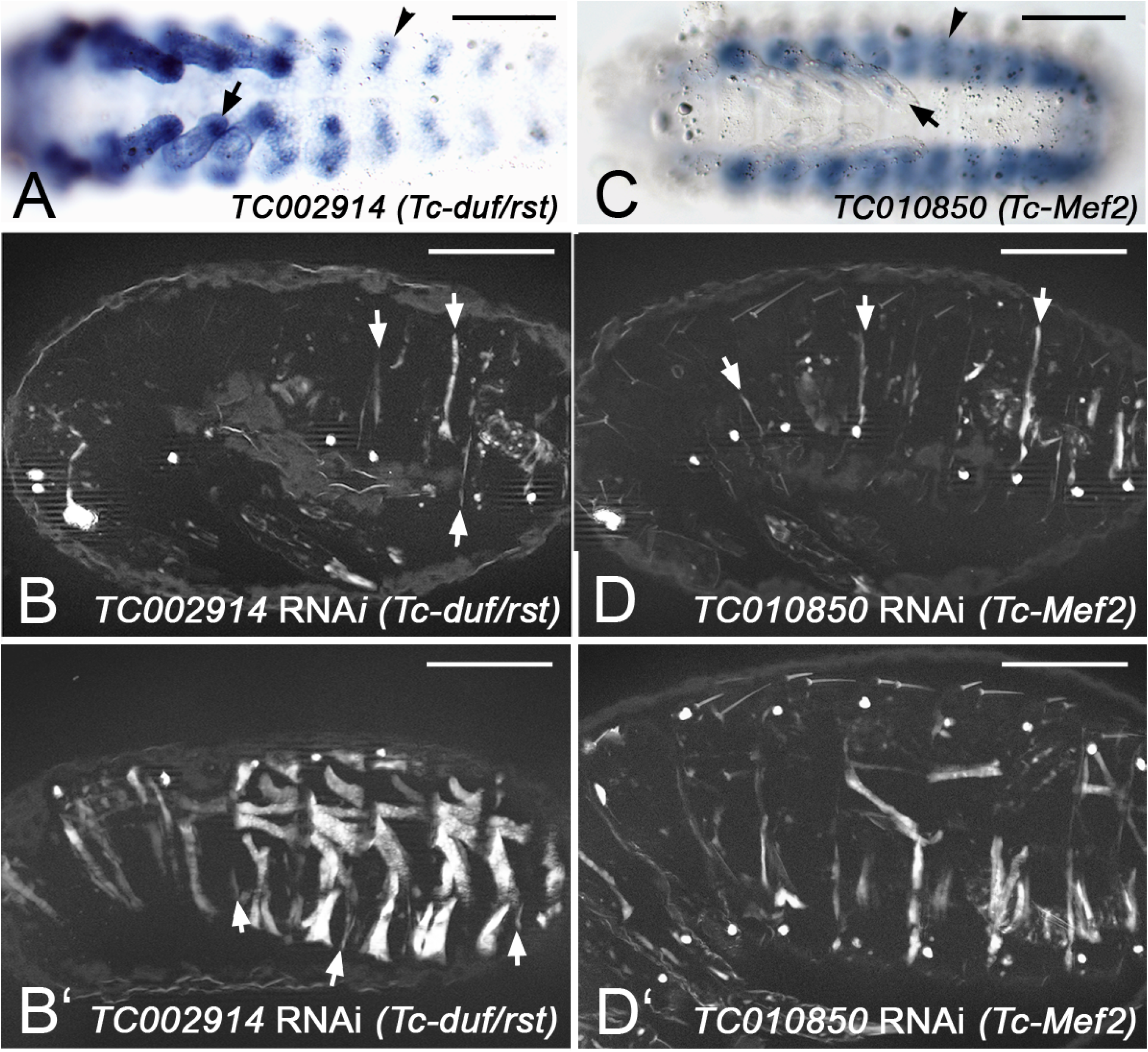
Expression and RNAi phenotypes of examples of *Tribolium* orthologs of known regulators of *Drosophila* muscle development. **(A)** *In situ* hybridization of *Tribolium* embryo (early germ band retraction stage) for mRNA of *TC002914* (ortholog of *Drosophila duf*/*rst*). Arrowhead: somatic mesoderm in abdominal segment. Arrow: Somatic mesoderm in leg. **(B)** Late stage *pig-19* embryo from adult female injected with dsRNA for *TC002914*, imaged live for EGFP. Only few and very thin muscle fibers are present (arrows). **(B’)** Example of milder phenotype in *TC002914* knock-down embryo than in (B), showing residual large muscles, very thin muscles (arrows), and gaps where muscles are missing. **(C)** *In situ* hybridization of *Tribolium* embryo (retracted germ band stage) for mRNA of *TC010850* (ortholog of *Drosophila Mef2*). Arrowhead: somatic mesoderm in abdominal segment. Arrow: Somatic mesoderm in leg. **(D)** Late stage *pig-19* embryo from adult female injected with dsRNA for *TC010850*, imaged live for EGFP. Few muscles are present that are very thin (arrows). **(D’)** Example of milder phenotype in *TC010850* knock-down embryo than in (D). Scale bars 100 m.

### First-pass pupal dsRNA injection screen to identify regulators of larval somatic muscle development

The main screen (‘pupal injection screen’), which we focus upon herein, involved injections of dsRNAs (named iB RNAs) for a total of ~ 8,500 genes into the body walls of female *pig-19* pupae. After crossing the eclosed females (if viable and fertile) with *black* males the EGFP-marked muscle patterns of their offspring were analyzed live under a compound fluorescence microscope in late embryonic stages and newly hatched first instar larvae, as shown for controls in Fig. 1. The screen was performed in two consecutive rounds and included screenings for various additional phenotypes by the screening consortium, as was described in an overview of the results from the first screening round of ~5,300 genes (Schmitt-Engel *et al.* 2015).

In the two rounds of first-pass screenings with pupal dsRNA injections, 205 of the ~8,500 tested genes were annotated with specific embryonic muscle phenotypes upon iB dsRNA injections that were not deemed to be secondary to broader disruptions such as segmentation defects, severe embryonic malformations, early developmental arrest, etc.. The muscle phenotypes were classified into broad categories and were annotated in the iBeetle database together with representative images. The classes with the largest numbers of representatives were those with missing muscles, with muscles featuring altered shapes, and with rounded and detached muscles, whereas those with incorrect muscle orientation and with specific effects on leg muscles were observed less frequently (Fig. 3A; Table S1). Particularly in the first round of first-pass screening, it turned out that primarily the classes “muscles missing” and “mainly/only leg muscles affected” contained many false-positives that were due to EGFP leakage from muscles upon injury of the tissues during preparation and mounting. In the second round of first-pass screening this effect was taken into account, which explains the smaller fractions of these two categories in this round (Fig. 3A).

**Figure 3.**
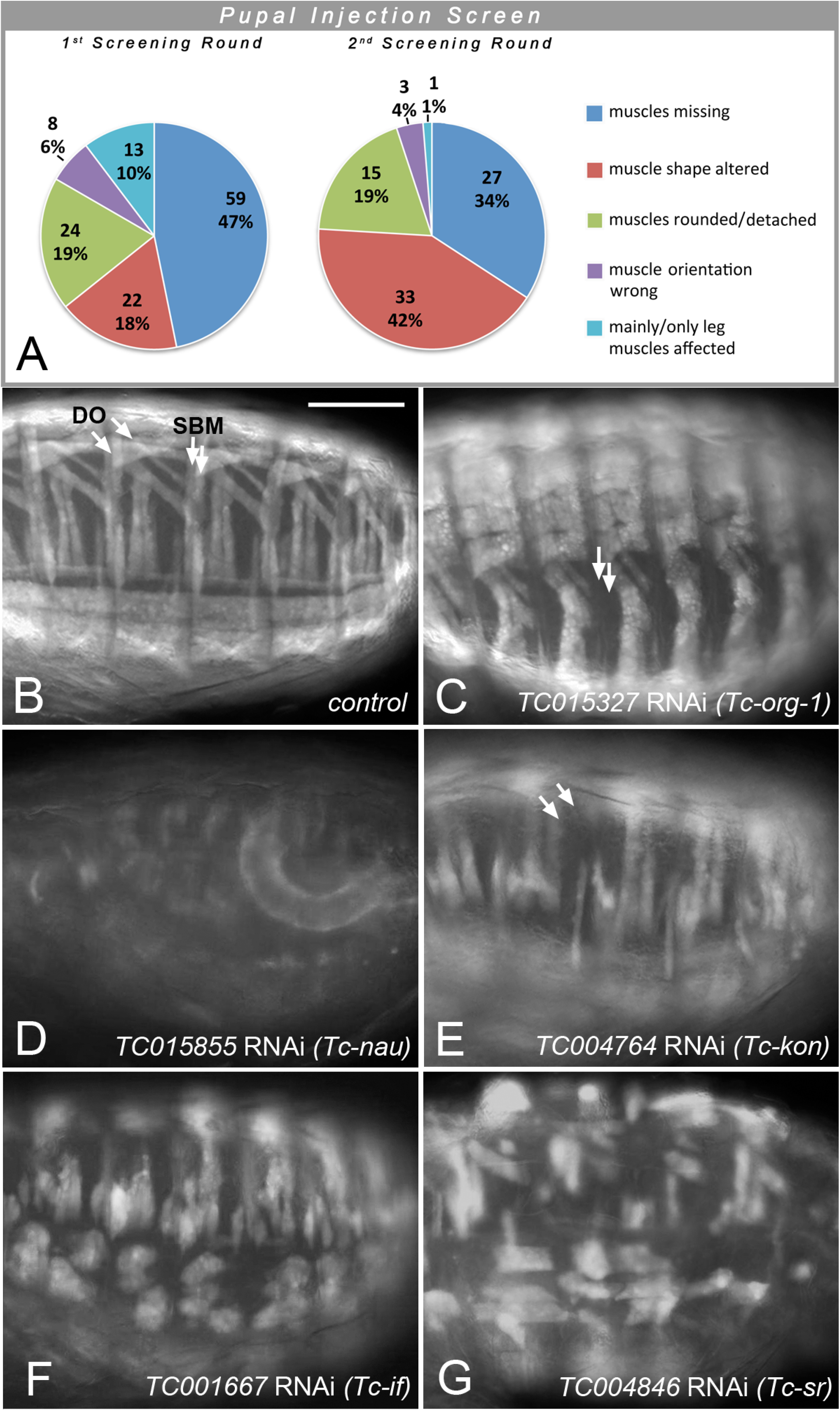
Classification and examples of larval muscle phenotypes obtained in the first-pass pupal injection screen. **(A)** The classification of larval muscle phenotypes of the total of 205 gene knock-downs annotated in the 1^st^ and 2^nd^ screening rounds are shown separately as the annotations in the categories “muscles missing” and “mainly/only leg muscles affected” from the first round included some technical artifacts that were largely circumvented in the 2^nd^ round. **(B)** Abdominal EGFP-marked muscles of *pig-19* late stage embryo (see Fig. 1E) from uninjected female shown as a control. DO: Dorsal oblique muscles; SBM: segment border muscles. **(C)** Embryo with RNAi knock-down of *TC015327* (*Tc-org-1*). Arrows indicate areas of missing or strongly reduced SBM muscles. **(D)** Embryo with RNAi knock-down of *TC015855* (*Tc-nau/MyoD*). **(E)** Embryo with RNAi knock-down of *TC004764* (*Tc-kon*). Arrows indicate areas of missing DO muscles. **(F)** Embryo with RNAi knock-down of *TC001667* (*Tc-if*) with detached and rounded muscles. **(G)** Embryo with RNAi knock-down of *TC004846* (*Tc-sr*) with detached and rounded muscles. Scale bar in (B) and also applicable to (C) - (G): 100 m.

As shown in Table 1, 24 of the 205 genes with annotated muscle phenotypes in the pupal injection screen corresponded to genes with *Drosophila* orthologs that have been implicated in various aspects of *Drosophila* muscle development. Although in this first-pass screen most of the muscle phenotypes were not characterized and annotated in detail, the phenotypes for the knock-downs of several *Tribolium* genes were reminiscent of the muscle phenotypes of mutations in their orthologs in *Drosophila*. For example, knock-down of the *Tribolium* ortholog of the *Drosophila* muscle identity gene *org-1* led to the absence of specific muscles, including the segment border muscles like in *Drosophila* (Fig. 3C, cf. Fig. 3B) (Schaub *et al.* 2012). Likewise, knock-down of the ortholog of *nautilus* (*nau*; *Drosophila MyoD*) led to a loss of muscles and reduction of the EGFP differentiation marker, although the observed phenotype appears more severe as compared to *Drosophila nau* mutants (Balagopalan *et al.* 2001) (Fig. 3D). Also knock-down of the *kon-tiki* (*kon*) ortholog caused the absence of subsets of muscle fibers, which in *Drosophila kon* mutants is attributed to defects in myotube migration and attachments in subsets of muscles (Schnorrer *et al.* 2007) (Fig. 3E). Knock-downs of the *Tribolium* orthologs of *inflated* (*if*) (Fig. 3F) and *stripe* (*sr*) (Fig. 3G) led to the appearance of spherical myotubes. This is likely due to disrupted muscle attachments because of weakened integrin-mediated adhesions with tendon cells (in the case of *if*) or the absence of differentiated tendon cells (in case of *sr*), as shown previously in *Drosophila* mutants of their respective orthologs (Brown 1994; Volk and VijayRaghavan 1994). The verification screens for novel candidates of myogenic regulators obtained in this first-pass screen and their RNAi phenotypes are described further below.

**Table 1.** **Compilation of screening data for *Tribolium* orthologs of *Drosophila* genes known to regulate various aspects of muscle development with clear *Tribolium* muscle phenotypes** (from primary screens; see also http://ibeetle-base.uni-goettingen.de)

| *Drosophila* Gene | Tc# | iB<br>dsRNA# | Pupal<br>Screen | Larval<br>Screen |
| --- | --- | --- | --- | --- |
|  |  |  | Muscle<br>phenotype | Muscle<br>phenotype |
| ***Ced-12*<sup>#</sup> (*Elmo*)** | TC002742 | 08003 | ✓ | --- |
| ***Crk*** | TC004767 | 07501 | ✓ | --- |
| ***dock*** | TC013803 | 02209 | **X** | ✓ |
| ***duf*<sup>#</sup> (*kirre*), *rst*<sup>#</sup>** | TC002914 | 03469 | **X**<sup>2)</sup> | ✓ |
| ***HLH54F*<sup>\*</sup>** | TC002349 | 06703 | ✓ | --- |
| ***htl*<sup>#</sup>** | TC004713 | 03821 | ✓ (no muscle-GFP) | **X**<sup>1)</sup> |
| ***if*<sup>#</sup>** | TC001667 | 03227 | ✓ | --- |
| ***kon*<sup>#</sup>** | TC004764 | 07523 | ✓ | --- |
| ***LanA*<sup>#</sup>** | TC003460 | 03560 | ✓ | ✓ |
| ***LanB1*<sup>#</sup>** | TC005184 | 08660 | ✓ | --- |
| ***LanB2*<sup>#</sup>** | TC010540 | 01705 | ✓ | **X** |
| ***lmd*<sup>#</sup>** | TC030749 | 06061 | ✓ | --- |
| ***nau*<sup>#</sup> (*MyoD*)** | TC015855 | 05915 | ✓ | --- |
| ***Ndg*** | TC008043 | 01306 | ✓ | **X** |
| ***numb*<sup>#</sup>** | TC012074 | 05163 | ✓ | **X** |
| ***org-1*<sup>\*</sup>** | TC015327 | 05796 | ✓ | --- |
| ***ptc*<sup>#</sup>** | TC004745 | 03831 | ✓ | **X**<sup>1)</sup> |
| ***salm*** | TC013501 | 10041 | ✓ | ---<sup>3)</sup> |
| ***simj*<sup>#</sup>** | TC009782 | 04697 | ✓ | **X** |
| ***siz*<sup>#</sup>** | TC005167 | 03913 | ✓ | **X** |
| ***sli*<sup>#</sup>** | TC032257 | 02993 | ✓ | --- |
| ***sr*<sup>#</sup>** | TC004846 | 03857 | ✓ | **X**<sup>1)</sup> |
| ***stumps*<sup>#</sup> (*dof*)** | TC011323 | 02631 | ✓ | **X** |
| ***Syd*** | TC003186 | 02682 | ✓ | **X** |
| ***twi*<sup>#</sup>** | TC014598 | 09112 | ✓ | --- |
| ***vg*** | TC010897 | 04931 | **X** | ✓ |
| ***Vrp1*<sup>#</sup>** | TC012341 | 05218 | ✓ | --- |
| ***wb*<sup>#</sup>** | TC014773 | 05688 | **X**<sup>1)</sup> | ✓ |

✓ muscle phenotype annotated
**X** no muscle phenotype annotated
--- no dsRNA injected
<sup>1)</sup> not screenable as RNAi caused lethality prior to muscle development (pupal, adult, or early embryonic)
<sup>2)</sup> but strongly depleted musculature in experiments with injections into adult beetles (see Fig. 1)
<sup>3)</sup> see reported *Tribolium* muscle phenotype in Schönbauer et al., Nature 479, 406-9 (2011)
<sup>\*</sup> muscle identity genes
<sup>#</sup> genes used for estimating the recovery rate of the screen for identifying *Tribolium* muscle regulatory genes. The marked genes have been reported and judged to give sufficiently conspicuous embryonic muscle phenotypes in *Drosophila* zygotic or germline clone mutants so that they should be detectable under conditions similar to those used in our pupal RNAi injection screen. Among muscle founder genes only those are included that affect more than ~15% of the muscles in each segment.

### First-pass larval dsRNA injection screen to identify regulators of adult indirect flight muscle development

A parallel screen (‘larval injection screen’) involved injections of a total of ~5,000 iB dsRNAs into L6 stage larvae of the *D17* strain and the analysis of the EGFP-marked late pupal thoracic muscle patterns (as well as additional phenotypes), as described in (Schmitt-Engel *et al.* 2015). In the first-pass larval injection screen, 96 of the ~5000 tested genes were annotated with specific pupal muscle phenotypes. Only six of these genes were also found in the pupal screen. More than half of the phenotypes were classified in the category of “altered muscle shapes” (52). Significantly fewer examples were found in the categories “muscles missing” (18), “muscles rounded and detached” (12), “muscle number increased/muscles split” (9), and “muscle orientation incorrect” (5) (Fig. 4A; Table S1). The number of muscle regulatory genes known from *Drosophila* showing phenotypes in *Tribolium* late pupal muscles upon knock-downs in the larval injections screen was significantly lower as compared to the pupal injection screen (Table 1 and data not shown).

**Figure 4.**
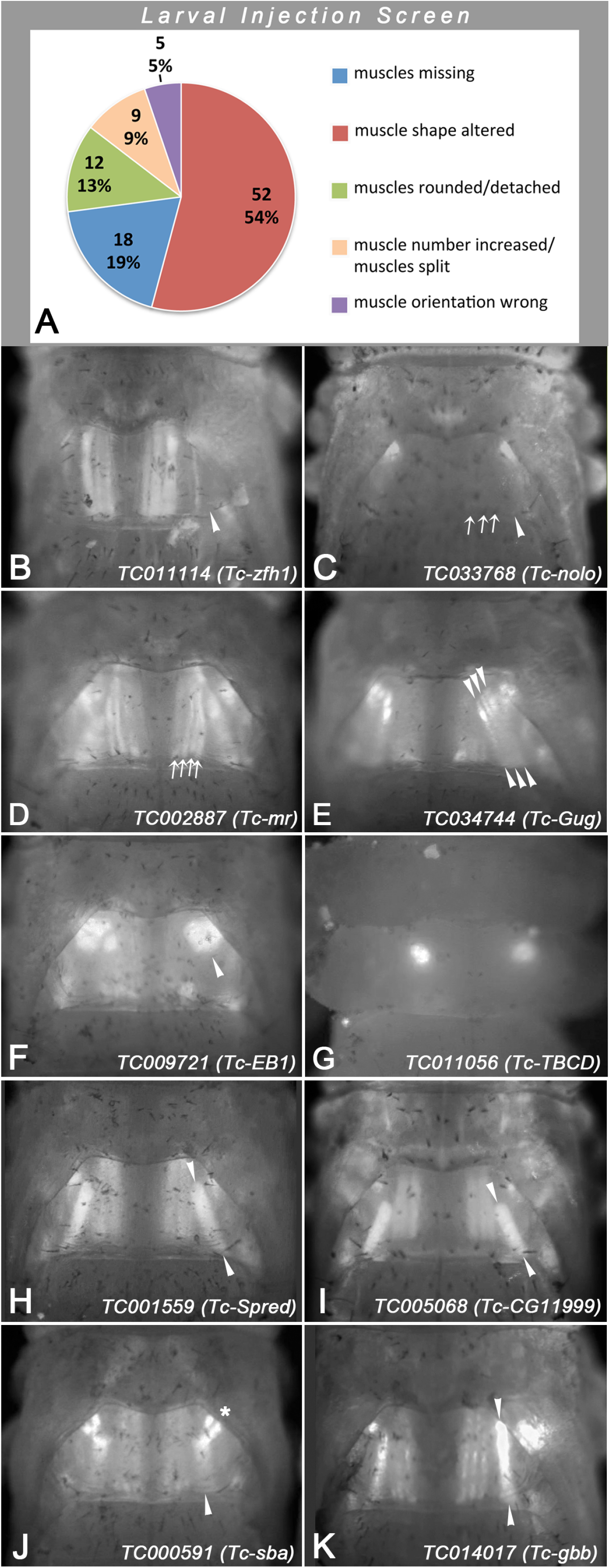
Classification and examples of pupal thoracic muscle phenotypes obtained in the first-pass larval injection screen. **(A)** Classification of pupal indirect flight muscle phenotypes of the total of 96 gene knock-downs annotated with such phenotypes in the first-pass larval injection screen. **(B - K)** Shown are dorsal views of thoraxes of *D17* pupae imaged and with magnifications as shown for control in Fig. 1K. Altered dorsal longitudinal flight muscles are indicated by arrows, dorsal oblique flight muscles by arrowheads, and dorsoventral flight muscles by asterisks in right halves of thoraxes. **(B)** Missing dorsal oblique muscles upon *TC011114* (*Tc-zfh1*) RNAi. **(C)** Undetectable dorsal longitudinal and dorsal oblique muscles upon *TC033768* (*Tc-nolo*) RNAi. **(D)** Increased number of dorsal longitudinal muscles upon *TC002887* (*Tc-mr*) RNAi. **(E)** Increased number of oblique muscles upon *TC034744* (*Tc-Gug)* RNAi. **(F)** Detached and rounded dorsal oblique muscles upon *TC009721* (*Tc-EB1*) RNAi. **(G)** Absence of muscles and presence of rounded syncytia upon *TC011056* (*Tc-TBCD*) RNAi. **(H)** Posteriorly shifted dorsal oblique muscle (particularly at its anterior insertion site) upon *TC001559* (*Tc-Spred*) RNAi. **(I)** Posteriorly shifted and shortened dorsal oblique muscle upon *TC005068* (*Tc-CG11999*) RNAi. **(J)** Mis-oriented dorsal oblique muscles and altered arrangements of dorsoventral muscles upon *TC000591* (*Tc-sba*) RNAi. **(K)** Mis-oriented dorsal oblique muscle and aberrant dorsoventral muscles upon *TC014017* (*Tc-gbb*) RNAi.

Fig. 4B - K shows representative examples of indirect flight muscle phenotypes from the first-pass larval injection screen (compare with control in Fig. 1J). Upon knock-down of *Tc-zfh1*, specifically the dorsal oblique flight muscles appear to be absent (Fig. 4B). With *Tc-nolo* RNAi, the dorso-longitudinal and dorsal oblique flight muscles are undetected (Fig. 4C). With *Tc-mr* RNAi, there are four instead of the normal three dorsal longitudinal flight muscles on either side (Fig. 4D). With *Tc-Gug* RNAi, there are three instead of a single dorsal oblique flight muscles present bilaterally (Fig. 4E). With *Tc-EB1* RNAi, particularly the dorsal oblique flight muscles are detached and rounded (Fig. 4F). With *Tc-TBCD* RNAi, the muscles are missing and instead a single rounded syncytium (or cluster of rounded syncytia) is present bilaterally (Fig. 4G). With *Tc-Spred* RNAi and *Tc-CG11999* RNAi, the dorsal oblique muscles are shifted posteriorly and, in the case of *Tc-CG11999* RNAi, it is shorter as compared to the control (Fig. 4H, I). With *Tc-sba* RNAi, the dorsal oblique muscles also appear shortened and oriented more longitudinally, and the dorsoventral flight muscles are arranged incorrectly as well (Fig. 4J). Likewise, with *Tc-gbb* the dorsal oblique muscles are arranged almost in parallel with the dorsal longitudinal muscles and the dorsoventral muscles display incorrect patterns (Fig. 4K). Unlike those from the pupal injection screen, the genes identified in the larval RNAi injection screen have not yet been subjected to verification screens to date.

The combined results from the first-pass pupal and larval screenings can be used for a rough estimation of the recovery rate of muscle regulatory genes. Among the 56 *Tribolium* orthologs of *Drosophila* genes connected with various roles in muscle development that were covered in the screens and could be evaluated, 27 showed a knock-down phenotype in late embryonic or (more rarely) in late pupal muscles (Table 1). For 29 genes no such phenotype was annotated in the first pass screens, 8 additional genes were not screenable for muscle phenotypes due to lethality, sterility, or broad embryo disruptions prior to muscle formation, and 14 have not been screened yet by dsRNA injections (Table S2). Under the (yet unproven) assumption that orthologous genes function similarly in muscle development of *Tribolium* and *Drosophila*, these comparisons would indicate a recovery rate of almost 50% of the muscle regulatory genes from *Tribolium* in our screens. If we look at the yields of the more complete pupal injection screen and select *Drosophila* genes with a very conspicuous muscle phenotype that should be recognizable under our screening conditions (if analogous), the recovery rate would be 56% (20 out of 36) (Fig. S1, Table 1, Table S2).

All these data, as well as the data for muscle phenotypes upon knock-downs of genes previously not implicated in muscle development can be accessed in the searchable iBeetle database (http://ibeetle-base.uni-goettingen.de; see listed iB numbers of the injected dsRNAs in Table S1), along with other morphological defects that were screened for in these large-scale RNAi screens (Dönitz *et al.* 2015; Dönitz *et al.* 2018).

### Verification screens of candidates for larval muscle regulators identified in pupal injection screen

In the next step, we performed a rescreen with 102 of the 229 iB dsRNAs that had annotated muscle phenotypes in the first-pass pupal injection screen in order to confirm these phenotypes (see Materials & Methods). In this rescreen, again performed in *pig-19*, the muscle phenotypes were confirmed for 54 of the original iB dsRNAs (Figure 5). As described above, particularly in the first round of first pass screening, many of the false-positives were due to EGFP leakage from muscles upon mechanical injury, and these were among the ones that got eliminated in this rescreen.

**Figure 5.**
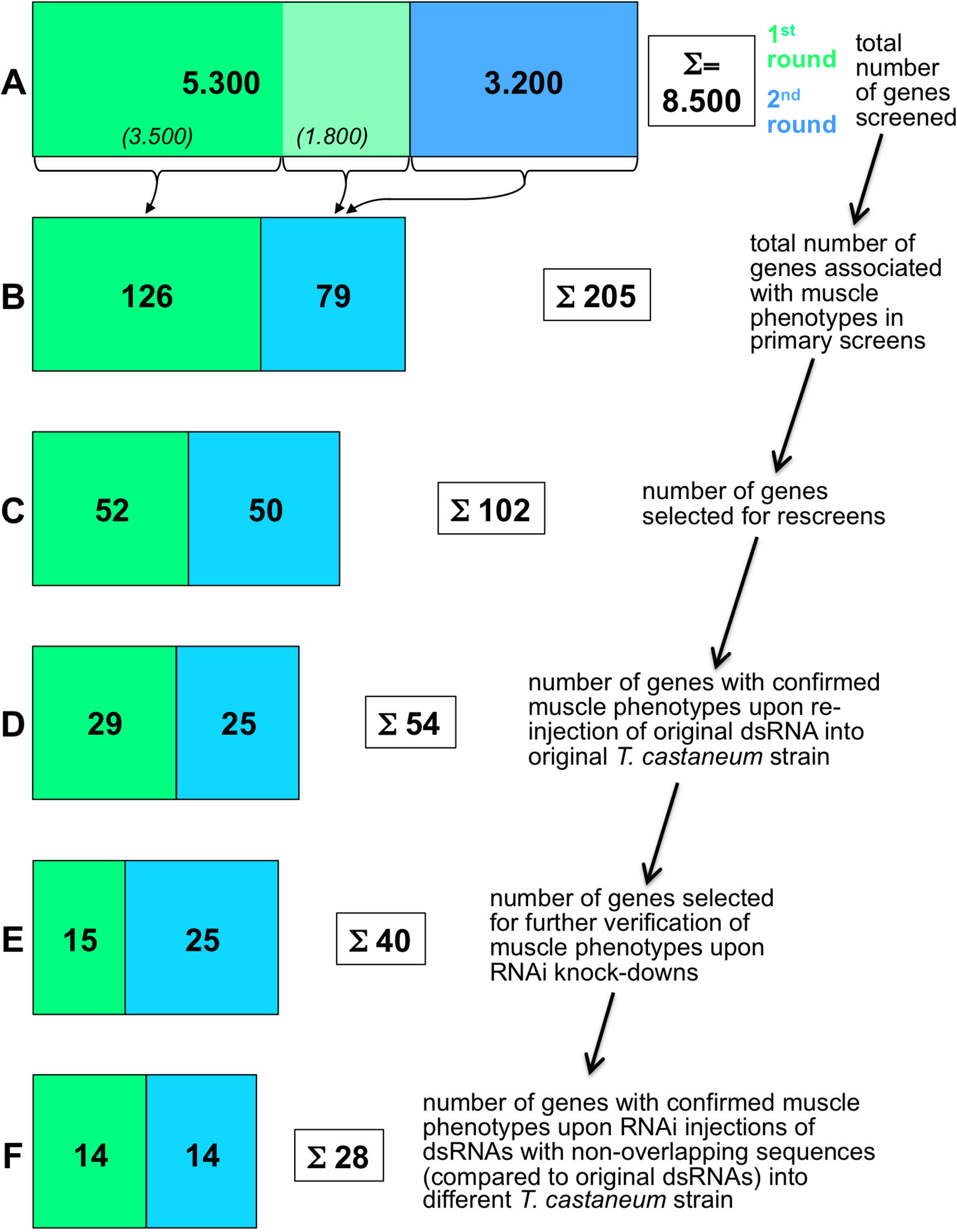
Progression of iBeetle RNAi screen for genes with knock-down phenotypes in larval muscles upon dsRNA injections into *Tribolium pig-19* pupae. Green areas in bars show numbers of screened and selected genes in the first round and blue areas the corresponding numbers in the second round of screening (blue-green area in (B) to (F) include “stragglers” from the first-round) (logarithmic scale). **(A)** A total of 8.500 genes were screened by dsRNA (“iB fragment”) injections in the two rounds. **(B)** 126 out of 3.500 injected dsRNAs from the first round and 79 out of another 1.800 injected dsRNAs from the first round and from 3.200 of the second round were annotated with muscle phenotypes in the primary screens. **(C)** 102 from the 205 genes in (B) were selected for rescreens. **(D)** 54 of the 102 genes from (C) were confirmed for embryonic muscle phenotypes upon re-injection of the original iB dsRNA fragments into *pig-19* pupae. **(E)** 40 of the 54 genes from (D) were selected for independent verification of the observed muscle phenotypes. **(F)** 28 of the 40 genes from (E) showed confirmed muscle knock-down phenotypes upon pupal injections of dsRNAs non-overlapping with the original iB dsRNA fragments (“NOFs”) into a different *T. castaneum* strain, *San Bernardino* (*SB*).

In a second rescreen, 40 of the corresponding genes were tested again with the aim to exclude off-target effects and possible strain specific effects. To reduce the work load, 14 genes were omitted, including some that showed only maternal expression in *Drosophila* or others that encoded enzymes with potentially broader or “house-keeping” functions. These included for example *TC010977* and *TC002552* that encode an elongase of very long fatty acids and a cytochrome P450, respectively. However we note that genes of this type may still have interesting functions in muscle tissues, see (Wang *et al.* 2016; Xu *et al.* 2018). For the 40 selected genes, new dsRNAs with sequences that did not overlap with the original iB dsRNA sequences were injected into female pupae from the San Bernardino (SB) strain, and the muscle patterns were analyzed in embryos from a cross of these females with *pig-19* males. In addition, the original iB dsRNA fragments were tested by analogous SB pupa injections. These tests served to rule out off-target effects and to confirm an essential role in muscle formation in different genetic backgrounds. As a result, the knock-down phenotypes in the embryonic musculature were confirmed for 28 of the 40 genes with the non-overlapping dsRNAs (Fig. 5). All 28 showed similar phenotypes in both *Tribolium* strains. These phenotypes and additional information on the affected genes, including the molecular features of their orthologs, are compiled in Table 2. The verified genes show a broad spectrum of distinct muscle phenotypes. Twelve of them exhibited rounded and detached muscles upon RNAi knock-down, eight had missing muscles, 6 showed muscles with altered shapes, and one showed effects mainly in the leg muscles (Table 2, bottom). The encoded proteins belong to a variety of different protein classes, most frequently being signaling proteins, DNA, chromatin, or RNA binding factors, and metabolic regulators (Table 2, bottom). and individual examples of them will be discussed in more detail in the following section.

**Table 2.**
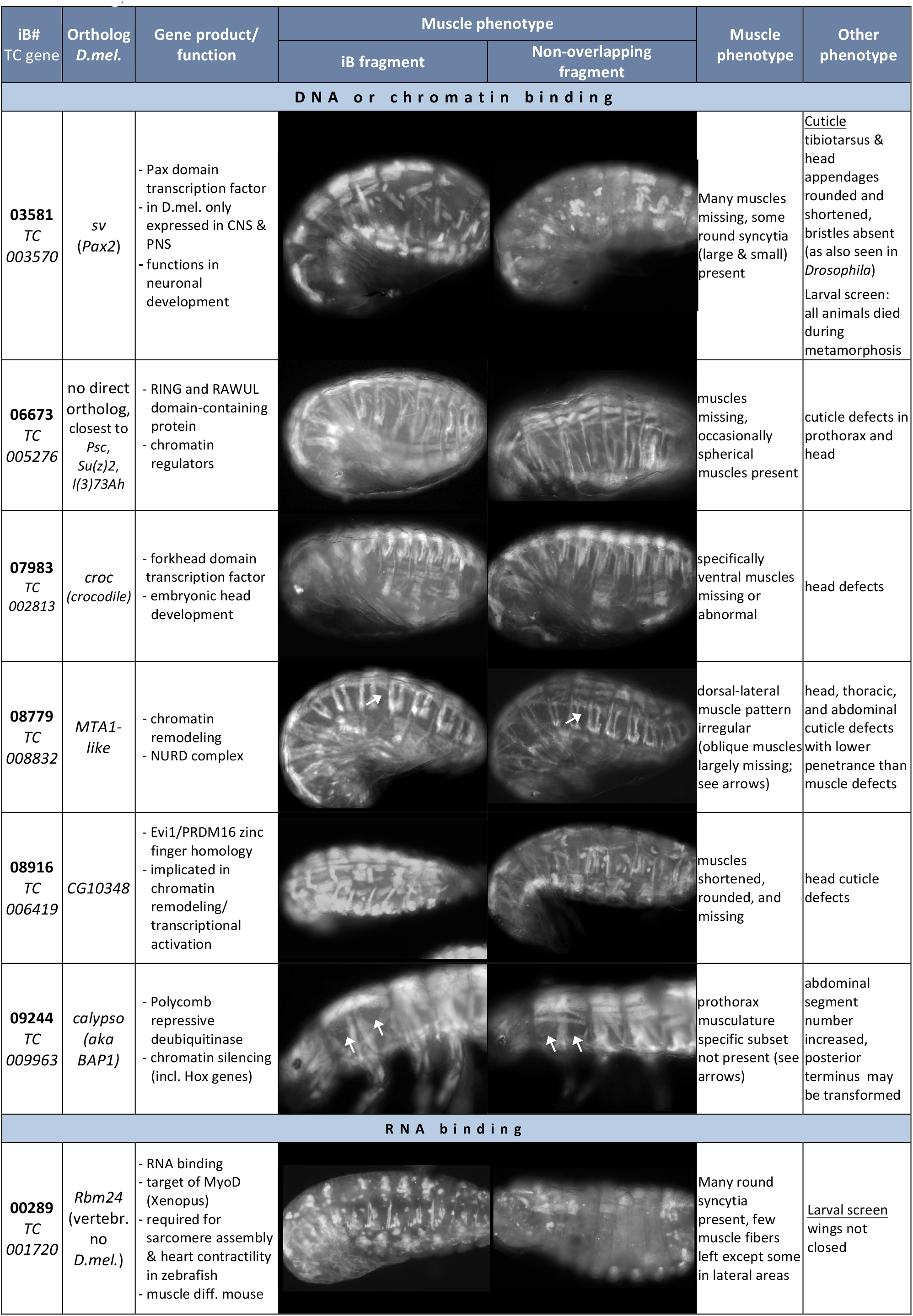
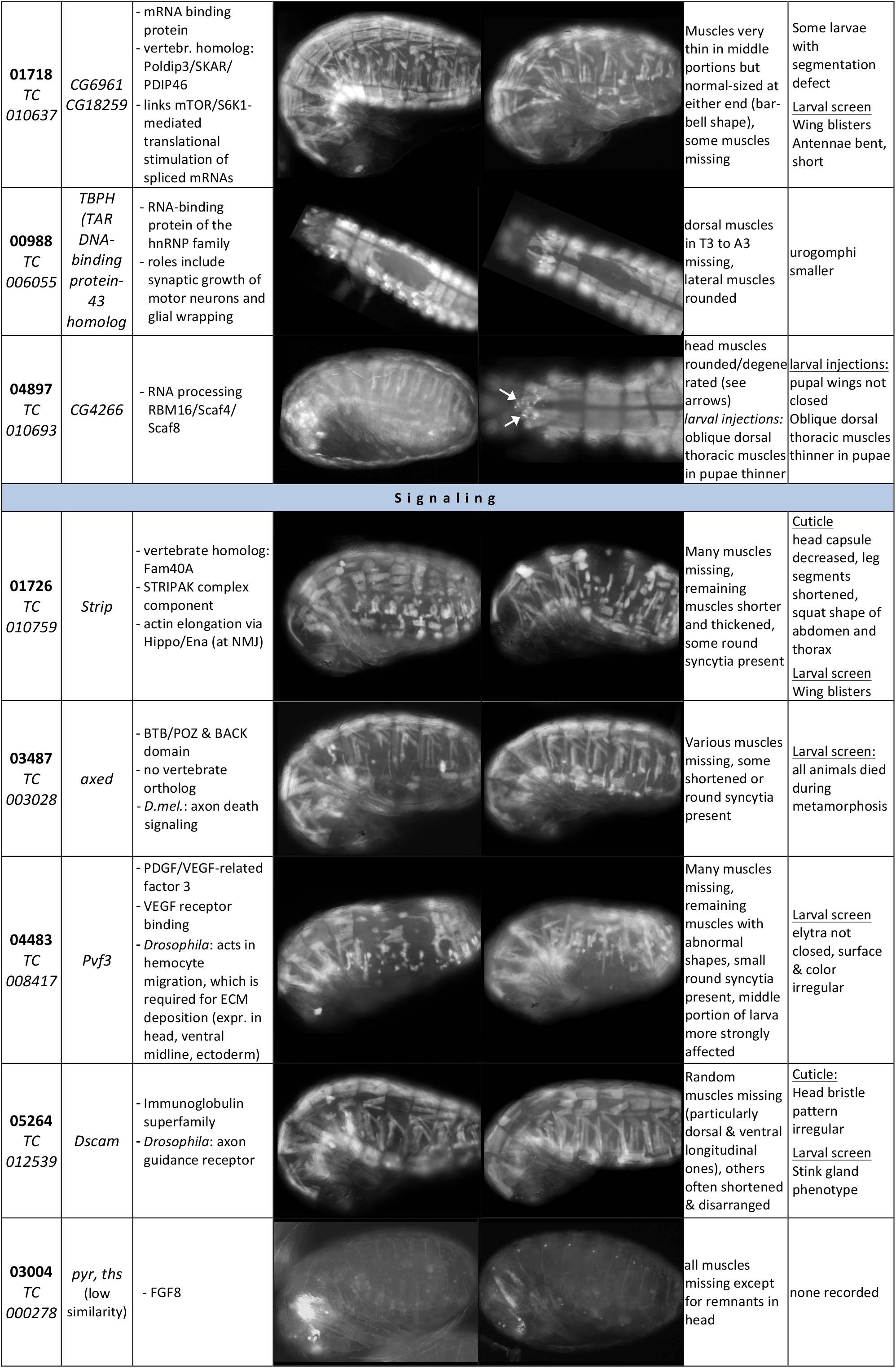
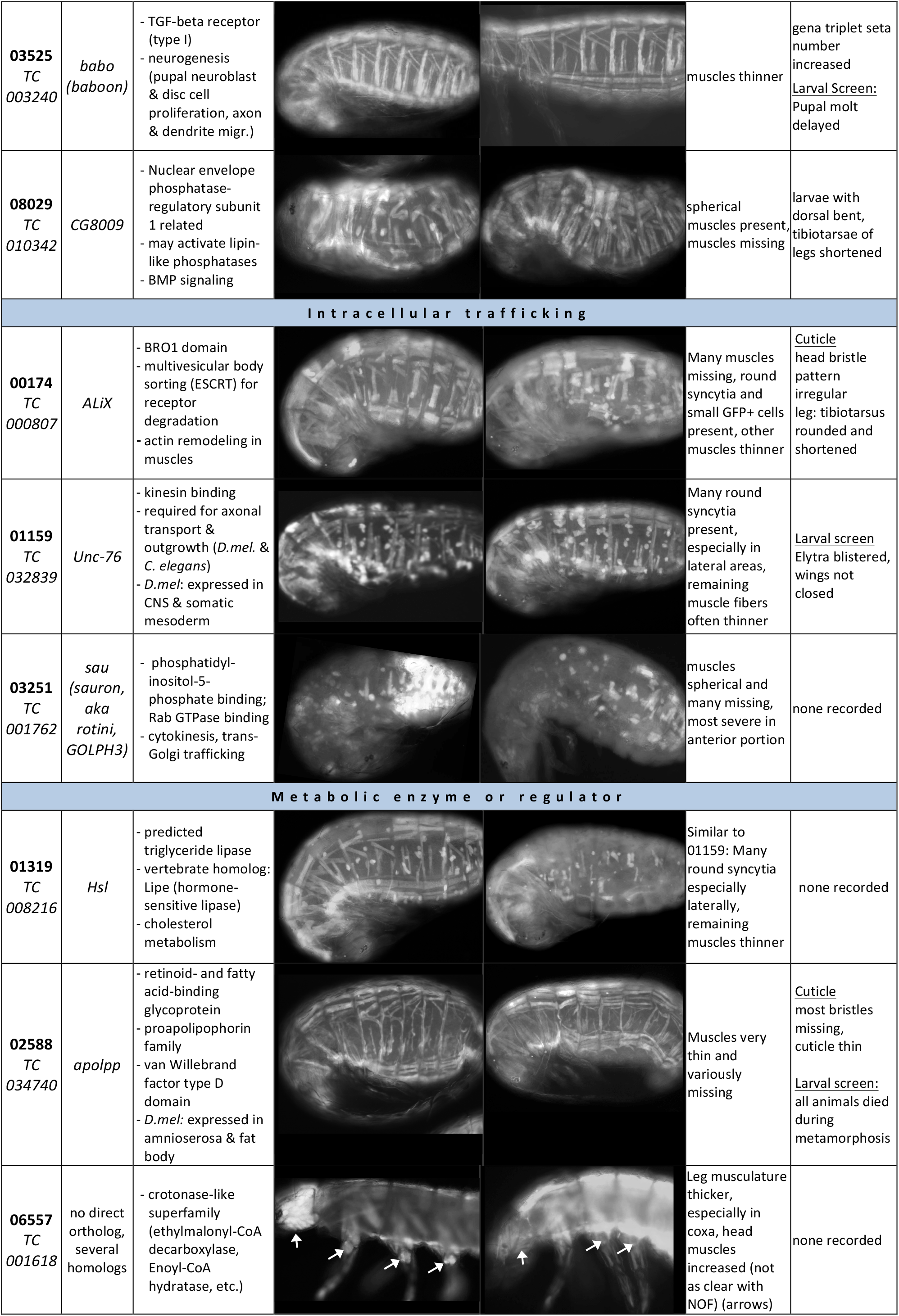
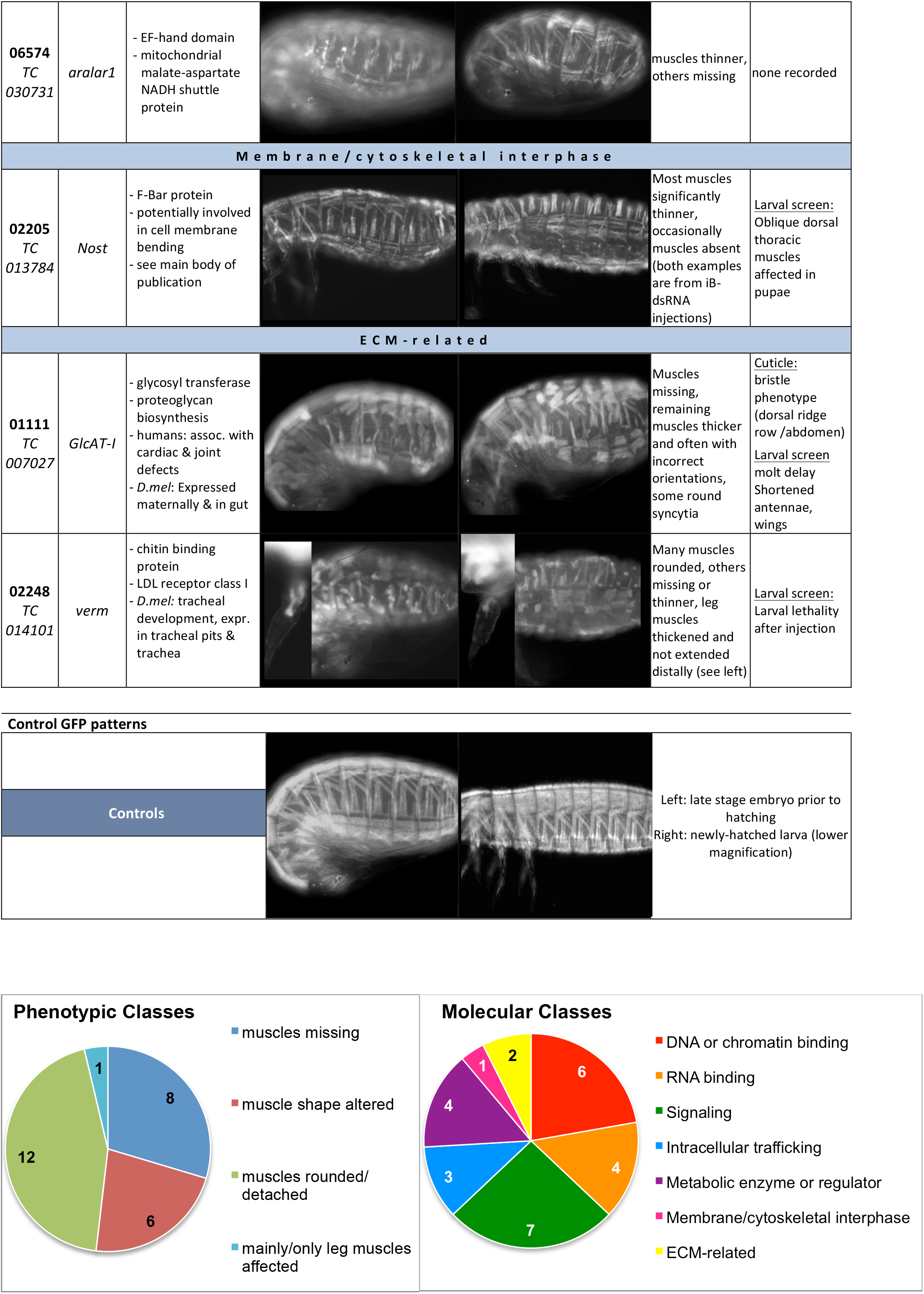
Features and verified larval muscle phenotypes of selected genes from the pupal injection screen. The phenotypes were confirmed in rescreens with original (iB) and non-overlapping (NOF) dsRNA fragments. Shown are representative images of the obtained phenotypes as well as controls (bottom). Genes are grouped by the molecular functions of the encoded proteins. The numerical distributions of classified phenotypes and molecular functions among these verified genes are shown at the end of the list.

| iB#<br>TC gene | Ortholog<br>*D.mel.* | Gene product/<br>function | Muscle phenotype |  | Muscle<br>phenotype | Other<br>phenotype |
| --- | --- | --- | --- | --- | --- | --- |
|  |  |  | iB fragment | Non-overlapping<br>fragment |  |  |
| DNA or chromatin binding |  |  |  |  |  |  |
| **03581**<br>TC<br>003570 | *sv*<br>(*Pax2*) | - Pax domain transcription factor<br>- in *D.mel.* only expressed in CNS & PNS<br>- functions in neuronal development |  |  | Many muscles missing, some round syncytia (large & small) present | <u>Cuticle</u><br>tibiotsarsus & head appendages rounded and shortened, bristles absent (as also seen in *Drosophila*)<br><u>Larval screen:</u><br>all animals died during metamorphosis |
| **06673**<br>TC<br>005276 | no direct ortholog, closest to *Psc*, *Su(z)2*, *l(3)73Ah* | - RING and RAWUL domain-containing protein<br>- chromatin regulators |  |  | muscles missing, occasionally spherical muscles present | cuticle defects in prothorax and head |
| **07983**<br>TC<br>002813 | *croc*<br>(*crocodile*) | - forkhead domain transcription factor<br>- embryonic head development |  |  | specifically ventral muscles missing or abnormal | head defects |
| **08779**<br>TC<br>008832 | *MTA1-like* | - chromatin remodeling<br>- NURD complex |  |  | dorsal-lateral muscle pattern irregular (oblique muscles largely missing; see arrows) | head, thoracic, and abdominal cuticle defects with lower penetrance than muscle defects |
| **08916**<br>TC<br>006419 | *CG10348* | - Evi1/PRDM16 zinc finger homology<br>- implicated in chromatin remodeling/transcriptional activation |  |  | muscles shortened, rounded, and missing | head cuticle defects |
| **09244**<br>TC<br>009963 | *calypso*<br>(*aka BAP1*) | - Polycomb repressive deubiquitinase<br>- chromatin silencing (incl. Hox genes) |  |  | prothorax musculature specific subset not present (see arrows) | abdominal segment number increased, posterior terminus may be transformed |
| RNA binding |  |  |  |  |  |  |
| **00289**<br>TC<br>001720 | *Rbm24*<br>(vertebr. no *D.mel.*) | - RNA binding<br>- target of MyoD (*Xenopus*)<br>- required for sarcomere assembly & heart contractility in zebrafish<br>- muscle diff. mouse |  |  | Many round syncytia present, few muscle fibers left except some in lateral areas | <u>Larval screen</u><br>wings not closed |

|  |  |  |  |  |  |
| --- | --- | --- | --- | --- | --- |
| **01718**<br>TC<br>010637 | *CG6961*<br>*CG18259* | - mRNA binding protein<br>- vertebr. homolog: Poldip3/SKAR/PDIP46<br>- links mTOR/S6K1-mediated translational stimulation of spliced mRNAs |  | Muscles very thin in middle portions but normal-sized at either end (barbell shape), some muscles missing | Some larvae with segmentation defect<br><u>Larval screen</u><br>Wing blisters<br>Antennae bent, short |
| **00988**<br>TC<br>006055 | *TBPH*<br>(*TAR* DNA-binding protein-43 homolog) | - RNA-binding protein of the hnRNP family<br>- roles include synaptic growth of motor neurons and glial wrapping |  | dorsal muscles in T3 to A3 missing, lateral muscles rounded | urogomphi smaller |
| **04897**<br>TC<br>010693 | *CG4266* | - RNA processing RBM16/Scaf4/Scaf8 |  | head muscles rounded/degenerated (see arrows)<br><u>larval injections:</u> oblique dorsal thoracic muscles in pupae thinner | <u>larval injections:</u> pupal wings not closed<br>Oblique dorsal thoracic muscles thinner in pupae |
| **Signaling** |  |  |  |  |  |
| **01726**<br>TC<br>010759 | *Strip* | - vertebrate homolog: Fam40A<br>- STRIPAK complex component<br>- actin elongation via Hippo/Ena (at NMJ) |  | Many muscles missing, remaining muscles shorter and thickened, some round syncytia present | <u>Cuticle</u><br>head capsule decreased, leg segments shortened, squat shape of abdomen and thorax<br><u>Larval screen</u><br>Wing blisters |
| **03487**<br>TC<br>003028 | *axed* | - BTB/POZ & BACK domain<br>- no vertebrate ortholog<br>- *D.mel.*: axon death signaling |  | Various muscles missing, some shortened or round syncytia present | <u>Larval screen:</u> all animals died during metamorphosis |
| **04483**<br>TC<br>008417 | *Pvf3* | - PDGF/VEGF-related factor 3<br>- VEGF receptor binding<br>- *Drosophila*: acts in hemocyte migration, which is required for ECM deposition (expr. in head, ventral midline, ectoderm) |  | Many muscles missing, remaining muscles with abnormal shapes, small round syncytia present, middle portion of larva more strongly affected | <u>Larval screen</u><br>elytra not closed, surface & color irregular |
| **05264**<br>TC<br>012539 | *Dscam* | - Immunoglobulin superfamily<br>- *Drosophila*: axon guidance receptor |  | Random muscles missing (particularly dorsal & ventral longitudinal ones), others often shortened & disarranged | <u>Cuticle:</u><br>Head bristle pattern irregular<br><u>Larval screen</u><br>Stink gland phenotype |
| **03004**<br>TC<br>000278 | *pyr, ths*<br>(low similarity) | - FGF8 |  | all muscles missing except for remnants in head | none recorded |
| **03525**<br>TC<br>003240 | *babo*<br>(*baboon*) | - TGF-beta receptor (type I)<br>- neurogenesis (pupal neuroblast & disc cell proliferation, axon & dendrite migr.) |  |  | muscles thinner | gena triplet seta number increased<br><u>Larval Screen:</u><br>Pupal molt delayed |
| **08029**<br>TC<br>010342 | CG8009 | - Nuclear envelope phosphatase-regulatory subunit 1 related<br>- may activate lipin-like phosphatases<br>- BMP signaling |  |  | spherical muscles present, muscles missing | larvae with dorsal bent, tibiotarsae of legs shortened |
| **Intracellular trafficking** |  |  |  |  |  |  |
| **00174**<br>TC<br>000807 | *ALiX* | - BRO1 domain<br>- multivesicular body sorting (ESCRT) for receptor degradation<br>- actin remodeling in muscles |  |  | Many muscles missing, round syncytia and small GFP+ cells present, other muscles thinner | Cuticle head bristle pattern irregular<br>leg: tibiotarsus rounded and shortened |
| **01159**<br>TC<br>032839 | *Unc-76* | - kinesin binding<br>- required for axonal transport & outgrowth (*D.mel.* & *C. elegans*)<br>- *D.mel.*: expressed in CNS & somatic mesoderm |  |  | Many round syncytia present, especially in lateral areas, remaining muscle fibers often thinner | <u>Larval screen</u><br>Elytra blistered, wings not closed |
| **03251**<br>TC<br>001762 | *sau*<br>(*sauron*,<br>*aka rotini*,<br>*GOLPH3*) | - phosphatidylinositol-5-phosphate binding; Rab GTPase binding<br>- cytokinesis, trans-Golgi trafficking |  |  | muscles spherical and many missing, most severe in anterior portion | none recorded |
| **Metabolic enzyme or regulator** |  |  |  |  |  |  |
| **01319**<br>TC<br>008216 | *Hsl* | - predicted triglyceride lipase<br>- vertebrate homolog: Lipe (hormone-sensitive lipase)<br>- cholesterol metabolism |  |  | Similar to 01159: Many round syncytia especially laterally, remaining muscles thinner | none recorded |
| **02588**<br>TC<br>034740 | *apolpp* | - retinoid- and fatty acid-binding glycoprotein<br>- proapolipoprotein family<br>- van Willebrand factor type D domain<br>- *D.mel.*: expressed in amnioserosa & fat body |  |  | Muscles very thin and variously missing | <u>Cuticle</u><br>most bristles missing, cuticle thin<br><u>Larval screen:</u><br>all animals died during metamorphosis |
| **06557**<br>TC<br>001618 | no direct ortholog, several homologs | - crotonase-like superfamily (ethylmalonyl-CoA decarboxylase, Enoyl-CoA hydratase, etc.) |  |  | Leg musculature thicker, especially in coxa, head muscles increased (not as clear with NOF) (arrows) | none recorded |

|  |  |  |  |  |  |
| --- | --- | --- | --- | --- | --- |
| 06574<br>TC<br>030731 | *aralar1* | - EF-hand domain<br>- mitochondrial malate-aspartate NADH shuttle protein |  | muscles thinner, others missing | none recorded |
| **Membrane / cytoskeletal interphase** |  |  |  |  |  |
| 02205<br>TC<br>013784 | *Nost* | - F-Bar protein<br>- potentially involved in cell membrane bending<br>- see main body of publication |  | Most muscles significantly thinner, occasionally muscles absent (both examples are from iB-dsRNA injections) | <u>Larval screen:</u><br>Oblique dorsal thoracic muscles affected in pupae |
| **ECM - related** |  |  |  |  |  |
| 01111<br>TC<br>007027 | *GlcAT-I* | - glycosyl transferase<br>- proteoglycan biosynthesis<br>- humans: assoc. with cardiac & joint defects<br>- *D.mel*: Expressed maternally & in gut |  | Muscles missing, remaining muscles thicker and often with incorrect orientations, some round syncytia | <u>Cuticle:</u><br>bristle phenotype (dorsal ridge row /abdomen)<br><u>Larval screen</u><br>molt delay<br>Shortened antennae, wings |
| 02248<br>TC<br>014101 | *verm* | - chitin binding protein<br>- LDL receptor class I<br>- *D.mel*: tracheal development, expr. in tracheal pits & trachea |  | Many muscles rounded, others missing or thinner, leg muscles thickened and not extended distally (see left) | <u>Larval screen:</u><br>Larval lethality after injection |

Control GFP patterns
|  |  |  |
| --- | --- | --- |
|  |  | Left: late stage embryo prior to hatching<br>Right: newly-hatched larva (lower magnification) |
| Controls |  |  |

## Discussion

### The iBeetle RNAi screen for muscle defects in *Tribolium* complements analogous screens in *Drosophila*

Using *Tribolium* as high-throughput RNAi screening platform, we uncovered a number of genes that had not been implicated in muscle development in previous work in *Drosophila*. Conversely, only about half of the orthologs of known *Drosophila* muscle genes (identified via both forward and reverse genetics) were recovered. This suggests that the different properties of these alternative screening platforms sometimes reveal different facets of a given biological process.

In *Drosophila*, several forward genetic screens using fluorescent reporter lines for somatic muscles have been performed. In two screens *MHC-tauGFP*, which marks all somatic muscles similar to the EGFP enhancer trap line used herein in *Tribolium*, was employed to screen for late embryonic muscle phenotypes upon ethyl methanesulfonate (EMS) mutagenesis (Chen and Olson 2001; Chen *et al.* 2008). Although the full screens have not been published, several mutants in genes regulating myoblast fusion, myotube targeting and attachment, and muscle maturation have been recovered from these (Chen and Olson 2001; Chen *et al.* 2003; Schnorrer *et al.* 2007; Johnson *et al.* 2013). Other EMS mutagenesis screens employed a cytoplasmic RFP reporter or a nuclear dsRed reporter driven by the founder cell enhancers of the muscle identity gene *org-1* and *apterous* (*ap*), respectively, in small subsets of muscles. From the *org-1*-RFP screen, mutants in the genes for the extracellular matrix proteins laminin β and collagen IV α1 have been reported to date, which revealed important roles of these proteins in muscle attachments, e.g., to the cardiac ECM (Hollfelder *et al.* 2014). From the ap::NLSdsRed screen, genes such as *esconsin* (*ens*) were identified that regulate nuclear positioning within myotubes (Metzger *et al.* 2012). For a number of reasons, including incomplete coverage of the genome and the possible failure to detect more subtle muscle phenotypes, none of these screens reached saturation. RNAi screens for muscle phenotypes upon gene knock-downs have been performed in *Drosophila* as well, which used lethality and locomotion or flight behavior as initial screening criteria. Instead of injections with dsRNAs, these screens employed inducible expression of dsRNAs with the UAS/GAL4 system and muscle-specific drivers. Due to the generally low knock-down efficiency of this method in embryonic stages, screening for phenotypes was largely confined to larval, pupal, and adult stages. In a large scale RNAi screen with *Mef2*-GAL4 driving transgenic UAS-IR RNA insertions, initial screening was for lethality and flightlessness, and follow-up analyses involved the examination of sarcomeric GFP markers (Schnorrer *et al.* 2010). This screen identified a large number of known and previously uncharacterized muscle-intrinsic players acting in muscle morphogenesis and function, including *spalt*, which turned out to be an evolutionarily-conserved master regulator of fibrillar flight muscle development during metamorphosis in insects (Schönbauer *et al.* 2011). Among the 23 genes with verified *Tribolium* muscle phenotypes (Table 2) that have *Drosophila* orthologs covered in the Schnorrer screen, only six were annotated for phenotypes in the primary screen in *Drosophila* (*CG11526*: early pupal lethality; *ths*: weak fliers; *sau*: semilethality; *babo*: late pupal lethality; *croc*: weak fliers; *MTA1-like*: flightlessness) and none of these were followed up with analyses of muscle phenotypes. Reasons for the viability and normal flight capabilities of the other 17 could include, 1) redundant gene functions in *Drosophila*, as exemplified by *Nostrin* in the accompanying paper (Schultheis *et al.* 2019), or mild defects not leading to lethality or overt flight defects; 2) diverged functions of the fly orthologs; 3) delayed functional knock-down in *Drosophila* embryos and absence of post-embryonic function in muscle development; 4) ineffective inverted repeat RNAs in *Drosophila* or false-positives in *Tribolium* in spite of the verification steps taken. In a recent small scale RNAi screen of 82 genes for larval locomotion defects, four genes with orthologs positive in the iBeetle screen were included: *twi* and *Vrp1*, which caused lethality, *if* which caused increased larval locomotion upon knock-down, and *nau* which lacked any locomotion phenotype. *singed* (*sn*, *Drosophila fascin*), which decreased locomotion, was shown to affect myoblast fusion to a similar degree as our *Nost cip4* double mutants (Camuglia *et al.* 2018; Schultheis *et al.* 2019). *Tc-sn* (*TC006673*) was not annotated with a muscle phenotype in the iBeetle screen.

In the iBeetle screens we exclusively used morphological defects as selection criteria in order to enrich for developmental regulators. We did not screen for locomotion defects or select for lethality that potentially could have yielded genes required for muscle physiology or function, such as those encoding sarcomeric proteins or their regulators. However, the percent lethality of the injected animals was annotated (Schmitt-Engel *et al.* 2015) and some, although not all knock-downs of genes for sarcomeric proteins covered by the screen did indeed yield high lethality of the injected animals within 11 days after pupal injections (e.g., 100% for *TC031441* [*Tc-TM1*]; 100% for *TC034001* [*Tc-MHC*], 60% for *TC001048* [*Tc-MLC1*]).

The iBeetle screen for genes with knock-down phenotypes in the somatic muscles had the technical advantage provided by the systemic agency of RNAi in *Tribolium* (Bucher *et al.* 2002), which made the injection work more economical due to the ease of injecting larvae and pupae and the recovery of large numbers of offspring from each injected female animal. In addition, this procedure is expected to knock down both the maternal and zygotic contributions of genes, and thus has the potential to uncover genes that may have been missed in the *Drosophila* screens due to maternal rescue. In contrast to the mesoderm-specific RNAi screens in *Drosophila* but similar to the EMS screens, the iBeetle RNAi screen involved global knock-downs of gene functions in all tissues of the animals. Hence, in addition to genes with muscle-intrinsic functions, genes with non-autonomous functions in muscle developments could be recovered as exemplified by *stripe* (Fig. 2), which in *Drosophila* is essential for muscle attachments through its function in determining epidermal tendon cell fates (Becker *et al.* 1997; Vorbrüggen and Jäckle 1997). On the other hand, global knock-downs increase the risk of secondary effects on muscle development, e.g., due to early effects of genes on developmental events prior to muscle development such as embryonic patterning, cell proliferation, etc.. These effects were minimized because the iBeetle screen included careful analyses of the larval cuticles with the aim to recover patterning genes as well (Schmitt-Engel *et al.* 2015; Ansari *et al.* 2018). Thus, even if some of the genes on our shortlist (Table 2) have additional roles in other tissues, their most prominent functions are expected to be in the development of the somatic musculature, either via mesoderm-intrinsic or via non-autonomous mechanisms.

As for any other screen, it is clear that we missed many genes affecting *Tribolium* muscle development. Among the *Tribolium* orthologs of 56 genes known to affect *Drosophila* muscle development that were included in the iBeetle screen, 27 were recovered through their knock-down phenotypes in the musculature. This is a surprisingly low portion given that the positive controls of genes with a variety of developmental functions included in the screen indicated a detection rate of about 80-90% (Schmitt-Engel *et al.* 2015). Only in a few cases this can be explained by the presence of two, potentially redundant *Tribolium* orthologs of genes that are represented only as single copies in *Drosophila* (such as *msh*, *Msp300*; Table S2). Further analyses are required to test whether the incomplete recovery of expected candidates is due to false negative annotations in the screen or to biological differences in the muscle developmental program between the two species of insect. One potential reason for missing larval muscle phenotypes in false-negatives could be the relatively long time delay between the pupal injections and the onset of embryonic muscle development, which in some cases may reduce the dsRNA concentrations below a critical threshold. As indicated by the rather severe phenotypes obtained with adult injections of *Tc-duf* dsRNAs as compared to those with pupal injections for several other myoblast fusion genes, adult injections may sometimes provide stronger effects. We did not systematically explore this possibility because pupal injections were necessary for other participants in the consortium, who screened for ovary and oogenesis defects (Schmitt-Engel *et al.* 2015). As in other screens with pan-muscle markers, it is also likely that some genes with more subtle knock-down phenotypes, e.g., affecting individual or small subsets of muscles, were missed and indeed, the majority of recovered phenotypes affected muscles globally (for examples of few exceptions, see *Tc-org-1*, Fig. 2C and *TC009963*, Table 2). This likely explains the low rate of identification of orthologs of *Drosophila* muscle identity genes, and accordingly, the recovery rate increases from 48% to 56% if the *Drosophila* muscle identity genes with phenotypes in only few muscles per hemisegment and other *Drosophila* genes with inconspicuous phenotypes are omitted (Table 1, Table S2).

### Novel genes identified in the iBeetle screen

Of note, many *Tribolium* genes were recovered for which their orthologs were not known to affect muscle development in *Drosophila*. A few genes lack any orthologs in *Drosophila* although some of them do have orthologs in vertebrates. A notable example for the latter is *Tc-RNA-binding motif protein 24* (*Rbm24)* (*TC001720*), for which knock-downs exhibited severe muscle phenotypes (Table 2; Schmitt-Engel *et al.* 2015). *Tc-Rbm24* mRNA is specifically expressed in the developing and mature embryonic somatic musculature and, more weakly, in the dorsal vessel (as well as in the CNS; DS and MF, data not shown). During our screen, mouse *Rbm24* (and presumably its paralog *Rbm38;* Miyamoto *et al.* 2009) was reported to play a major role in embryonic skeletal muscle development by regulating alternative splicing of a large number of muscle-specific primary transcripts (Yang *et al.* 2014). It is conceivable that in *Drosophila* the role of *Tc-Rbm24* in muscle development is exerted by other members of the RRM superfamily of RNA binding proteins. Starting to explore this possibility, we found that some *Drosophila* RRM superfamily members are expressed in the embryonic somatic mesoderm (*boule*, *CG33714*, *Hrb87F*; in the case of *boule* exclusively so; (Schultheis 2016). In addition to *Tc-Rbm24*, three other genes encoding putative *Tribolium* RNA binding proteins were recovered (*TC010637*, *TC006055*, *TC010693*; Table 2), reinforcing the important contribution of RNA metabolism in regulating normal muscle development (see also examples from *Drosophila* (Volk *et al.* 2008; Johnson *et al.* 2013; Oas *et al.* 2014; Spletter *et al.* 2015).

Several other identified genes shown in Table 2 encode predicted chromatin regulators, which are likely to influence gene regulatory programs during muscle development (*TC005276*, *TC005276*, *TC006419*, *TC009963*) and possibly have additional, perhaps less prominent roles in other tissues. *Tc-croc* (*TC002813*, Table 2), which encodes a forkhead domain transcription factor, exclusively affects the ventral muscles, suggesting that it may act as a muscle identity gene in *Tribolium*. Interestingly, *Drosophila croc* appears to be expressed in subsets of ventral mesodermal cells during early muscle development, but we have been unable to detect any ventral muscle defects in *Drosophila croc* mutant embryos (Häcker *et al.* 1995; MW and MF, unpublished data).

The *Drosophila* and mammalian counterparts of TC032839 protein, Unc-76 and FEZ1, respectively, bind to the Kinesin-1 Heavy Chain (KHC). FEZ1 binding in combination with JNK interacting proteins (JIP1, and perhaps similarly JIP3) was shown to release Kinesin-1 autoinhibition and thus activate the motor protein for microtubule binding and motility (Blasius *et al.* 2007; Koushika 2008). In accordance with this molecular interaction, *Drosophila* and *C. elegans unc-76* were shown to be required for axonal outgrowth and transport (Bloom and Horvitz 1997; Gindhart *et al.* 2003). In *Drosophila*, kinesin and kinesin-associated proteins, including JIP1/Aplip1 and JIP3/Synd, were shown to regulate nuclear positioning within muscle syncytia (Metzger *et al.* 2012; Schulman *et al.* 2014; Auld *et al.* 2018). Therefore we presume that Unc-76 likewise is involved in this regulatory pathway. *Tc-Unc-76* and *Dm-Unc-76* are both expressed in the somatic mesoderm (and more prominently in the CNS, as well as maternally), but unlike with RNAi in *Tribolium*, CRISPR/Cas9-generated zygotic null mutants did not show any overt muscle morphology phenotype in *Drosophila* embryos (DS, MW, and MF, unpublished data). Therefore, future analyses should investigate myonuclear positioning in *Drosophila* mutants that lack both the maternal and the zygotic contributions of *Unc-76*.

In addition, three signaling components were identified, namely Tc-fgf8, Tc-Babo, and Tc-Pvf3. *TC-fgf8* (*TC000278*) encodes the single *Tribolium* FGF8 member and is a putative ligand for the single, mesodermally-expressed FGF receptor Tc-fgfr, both of which were recently shown to be required for maintaining the expression of Tc-Twist in the somatic mesoderm of late stage embryos (Sharma *et al.* 2015). This requirement, along with a possible requirement for the activation of yet undefined differentiation genes, could explain the complete absence of *pig-19* muscle EGFP in *Tc-fgf8* RNAi embryos. It will be interesting to investigate in more detail how far muscles can develop in the absence of FGF8 signals. *Tc-babo* (*TC003240*) encodes a *Tribolium* TGF-beta type 1 receptor. The thin-muscle phenotype upon knock-down is reminiscent of phenotypes obtained upon knock-downs of myoblast fusion genes (Schultheis *et al.* 2019) and in this case could perhaps be due to under-proliferation of the fusion-competent myoblasts. *Tc-Pvf3* (*TC008417*) encodes a putative ligand for the *Tribolium* PDGF/VEGF related receptor. In *Drosophila* this ligand/receptor interaction is required for normal proliferation, migration, and maintenance of hemocytes (Parsons and Foley 2013; Sopko and Perrimon 2013). Hemocytes, in turn, are required for the deposition of extracellular matrix components in the *Drosophila* body wall (Matsubayashi *et al.* 2017). Thus it is conceivable that the phenotype of rounded muscles seen upon *Tc-Pvf3* knock-down is due to muscle detachments as a result of deficient extracellular matrix at their attachment sites (Maartens and Brown 2015). Following up on these avenues, and likewise on the exact involvement of the other genes identified in the screen, could yield unexpected insights into new aspects in the regulation of insect muscle development. In addition, verification screens for the candidates obtained in the larval injection screen should yield interesting new players during adult indirect flight muscle development. Additional preliminary characterizations of identified genes are available online in (Schultheis 2016) (in German), and a detailed examination of the *Drosophila* ortholog of one of them, *Nostrin*, is provided in the accompanying manuscript (Schultheis *et al.* 2019).

## Conclusion

The iBeetle RNAi screen has identified numerous genes that are candidates for regulators of insect muscle development. In several cases, their orthologs were already known to play roles in muscle development in *Drosophila*, and in some cases in vertebrates. A significant number of the genes identified in *Tribolium* had not been recovered in previous *Drosophila* work before. It will be interesting to examine the roles of identified genes previously not implicated in muscle development in detail in both *Tribolium* and in *Drosophila*. In *Tribolium*, the functional studies by RNAi can now be complemented by CRISPR/Cas9 induced mutations and engineered gene loci (Gilles *et al.* 2015). In parallel, more detailed analyses of the process of *Tribolium* muscle development will provide interesting insight into the similarities and differences of muscle development in beetle as compared to fly embryos. In addition, the *Drosophila* orthologs of newly identified genes with muscle phenotypes in *Tribolium* can be studied for potential functions in fly muscle development and added to the well-developed framework of regulatory networks in this system. The accompanying paper (Schultheis *et al.* 2019) presents an example of this approach by showing that the *Drosophila* ortholog of the F-Bar domain encoding gene *Tc-Nostrin*, identified in the iBeetle screen through its muscle phenotype, together with related F-Bar proteins plays a role in *Drosophila* myoblast fusion and the morphogenesis of adult midgut muscles. Functional redundancy in the fly previously had impeded the identification of this role in *Drosophila*.

## Acknowledgments

We acknowledge support by the German Research Foundation (DFG) for the *iBeetle* project (FOR1234). We thank Michael Schoppmeier and Martin Klingler for their vital contributions to the organization of the screening at Erlangen.

## Supplementary files

**Table S1.**
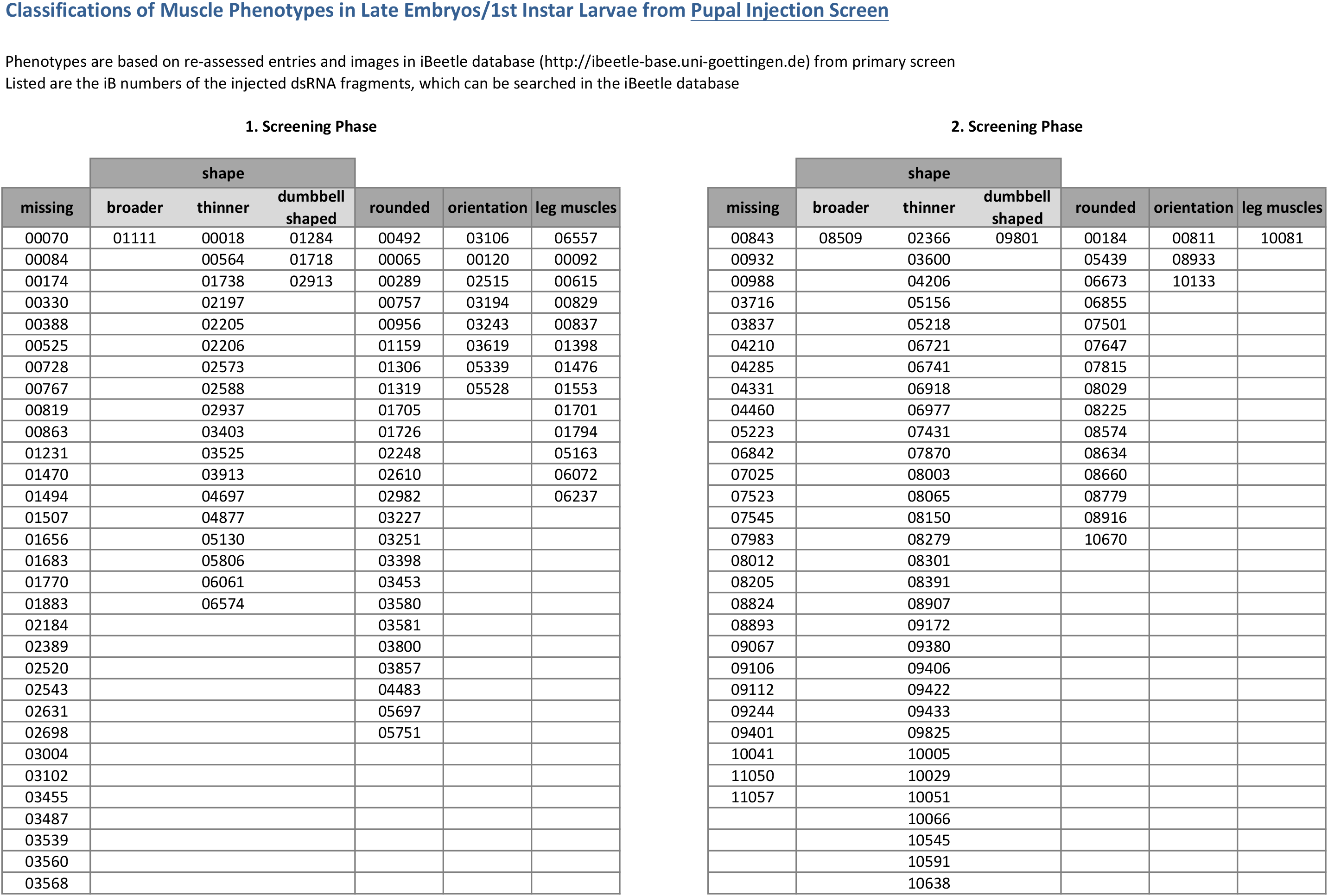
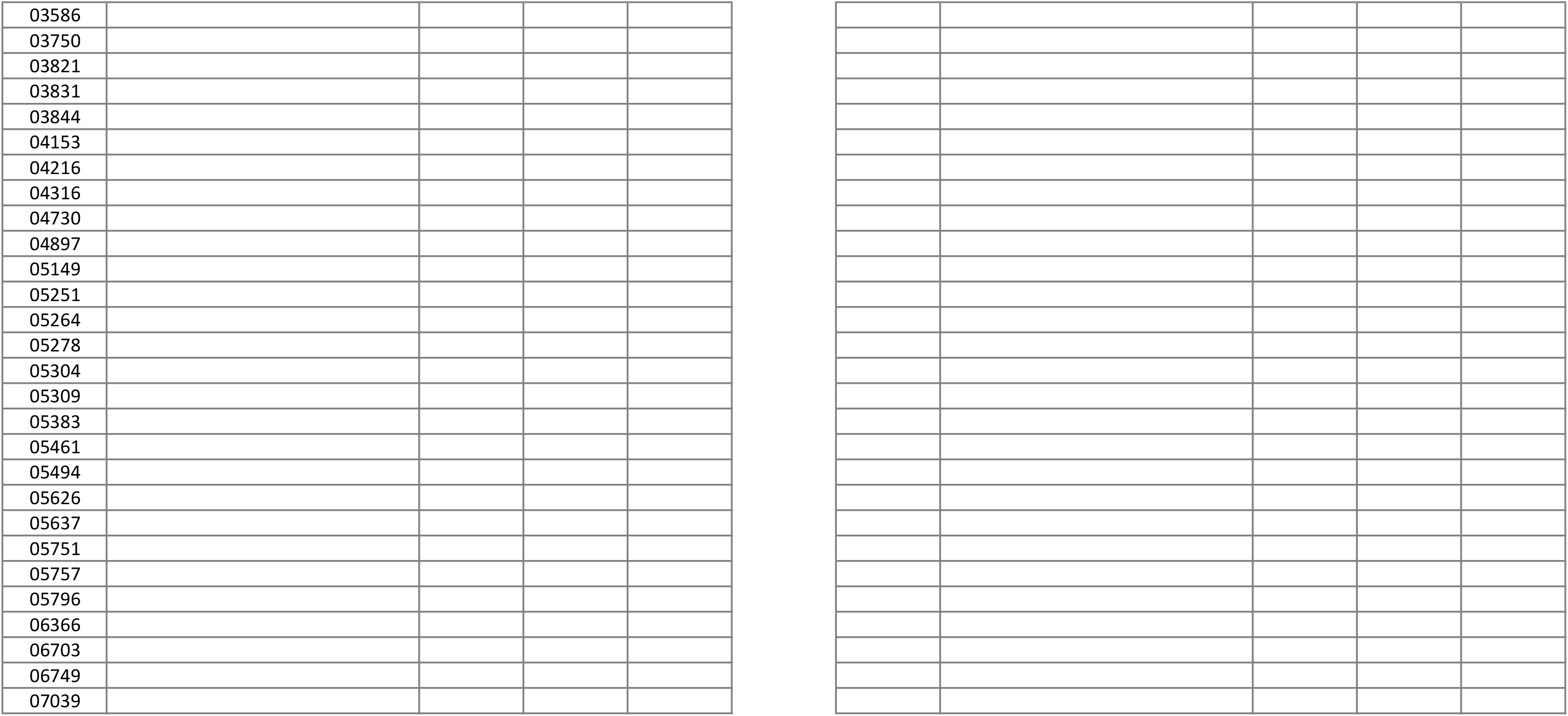
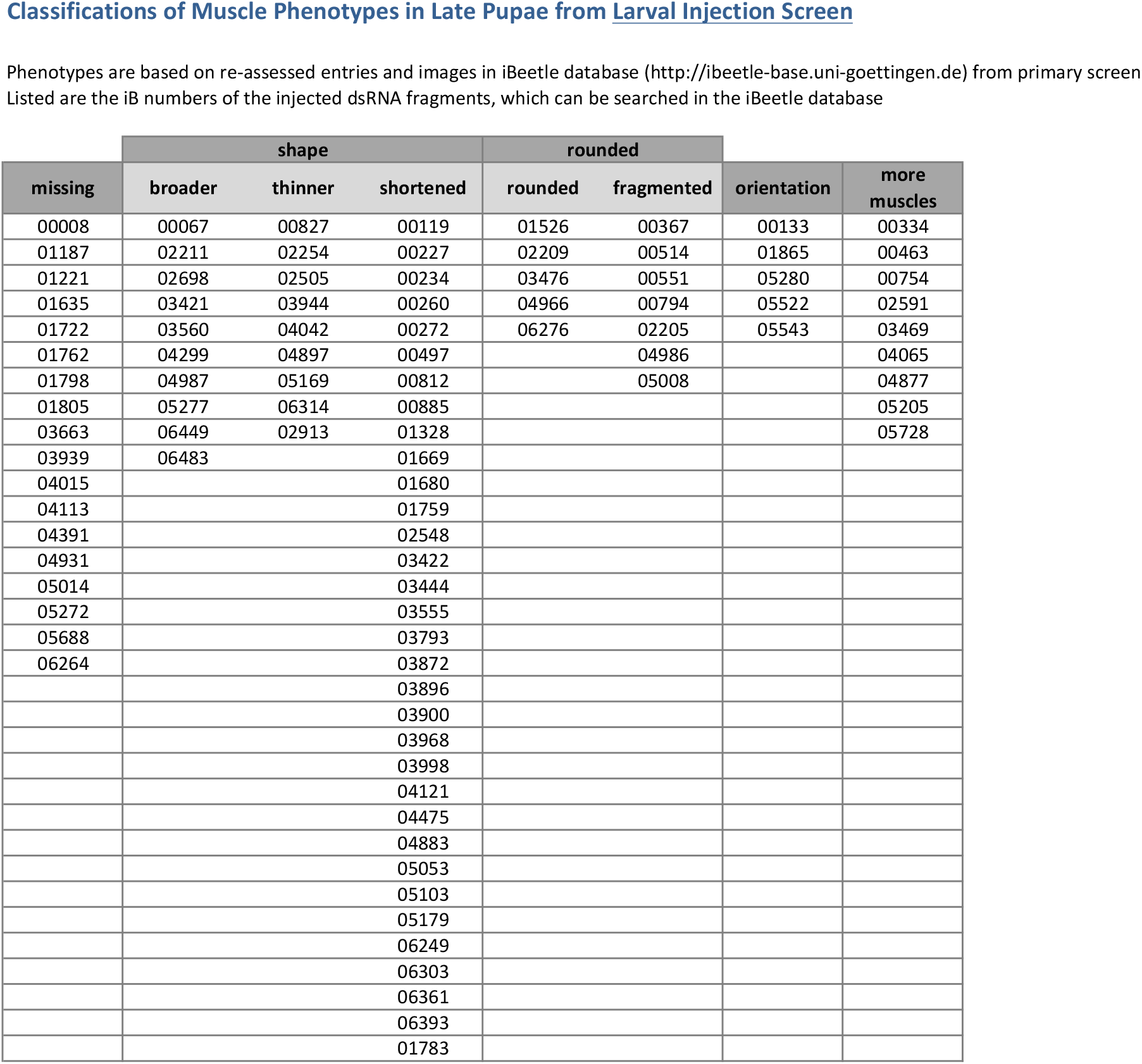
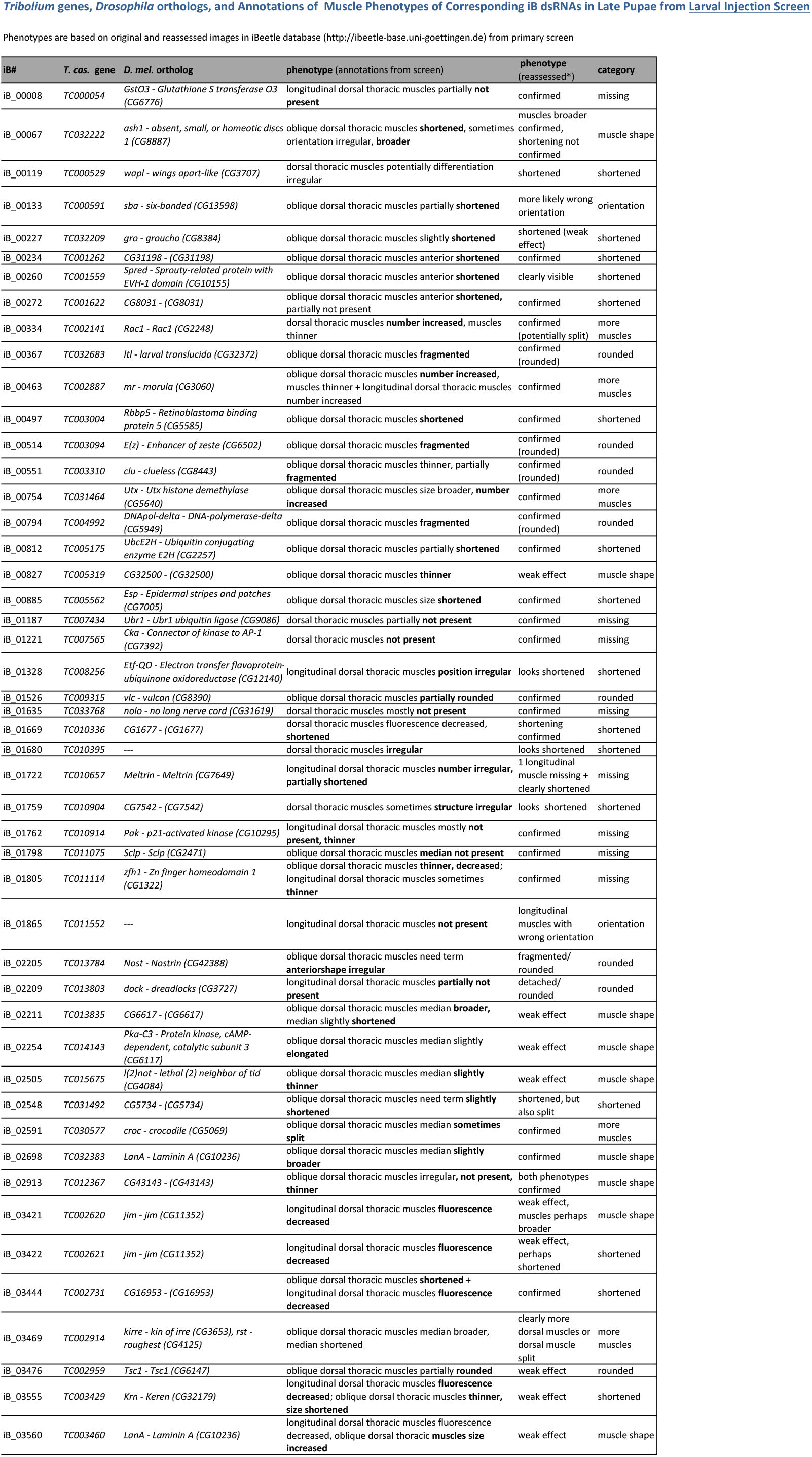
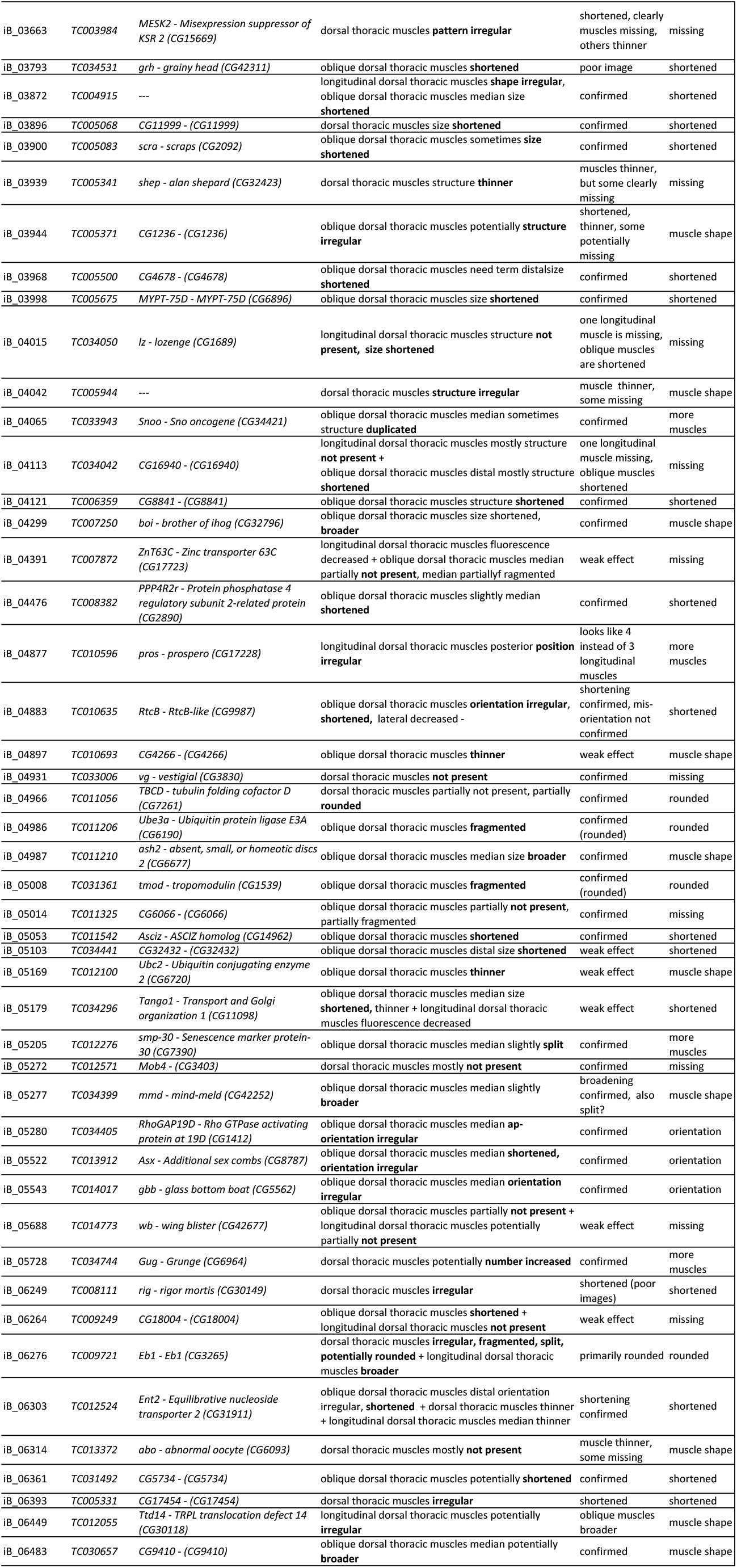
Lists of all dsRNA fragments with annotated muscle phenotypes from first-pass screens upon larval and pupal dsRNA injections. iB numbers denote the original dsRNA fragments used in primary screens and are grouped according to phenotypic classes. The numbers can be used to access full information on the respective phenotypes and corresponding genes in the iBeetle database (http://ibeetle-base.uni-goettingen.de, “Gene search”). The database entries were generated during the screen by the different screeners. Note that some of the final classifications were reassessed by a single expert (DS) post-screen but left unchanged in the database. For the larval injection screen additionally the *T. cas.* gene names, *D. mel.* orthologs, and the original as well as the reassessed phenotypes are listed. For the verified genes in the pupal screen these and additional data are included in Table 2.

**Table S2.**
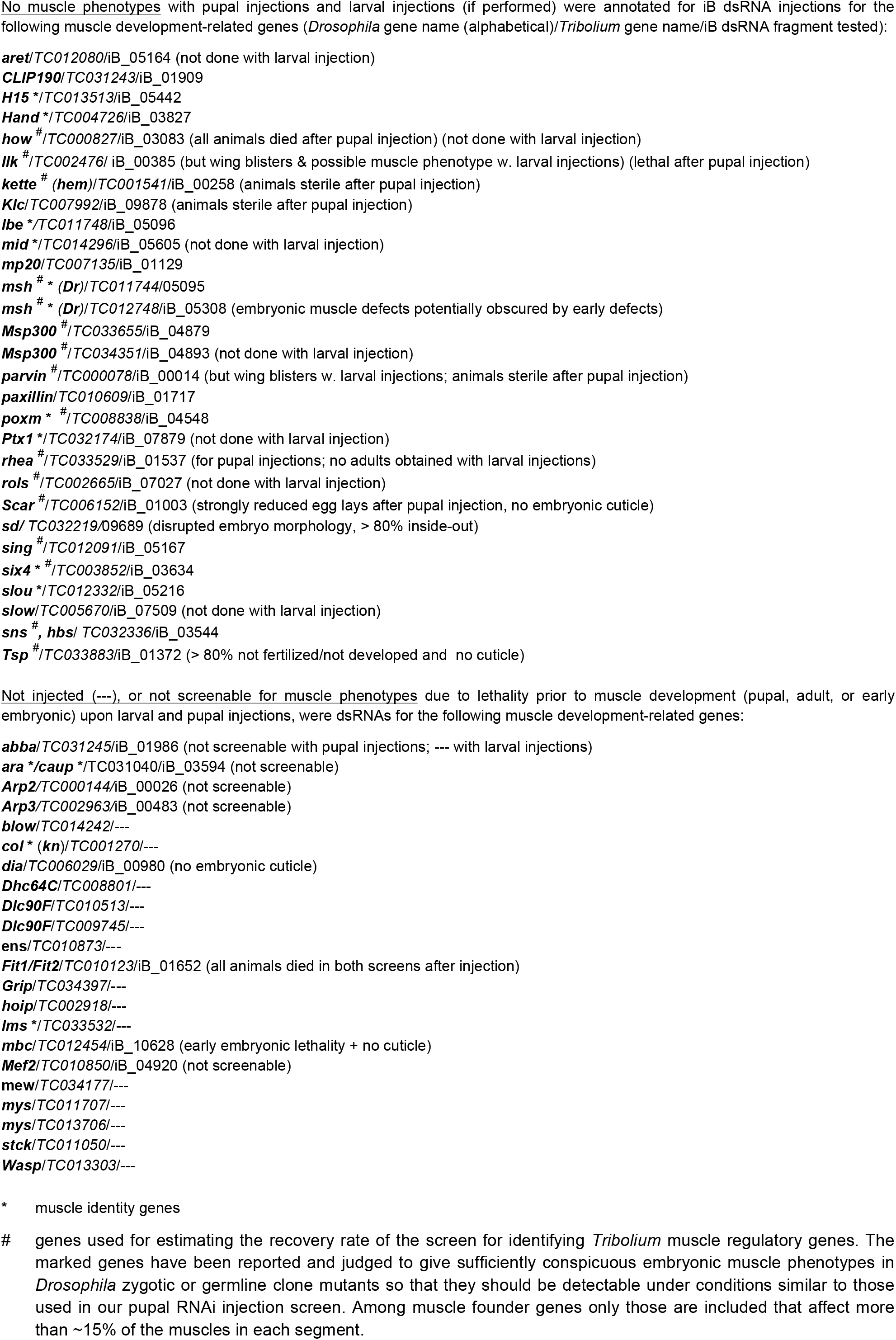
Compilation of screening data for *Tribolium* orthologs of *Drosophila* genes known to regulate various aspects of muscle development with no annotated *Tribolium* muscle phenotypes, uninterpretable muscle phenotypes, or no injected dsRNAs.

